# Memory CD8^+^ T cells mediate early pathogen-specific protection through localized delivery of chemokines and IFNγ to clusters of inflammatory monocytes

**DOI:** 10.1101/2021.03.01.433468

**Authors:** Marie Boutet, Zachary Benet, Erik Guillen, Caroline Koch, Saidi M’Homa Soudja, Fabien Delahaye, David Fooksman, Grégoire Lauvau

## Abstract

While cognate antigen drives clonal expansion of memory CD8^+^ T cells to achieve sterilizing immunity in immunized hosts, not much is known on how cognate antigen contributes to early mechanisms of protection before clonal expansion occurs. Herein, using distinct models of immunization, we establish that cognate antigen recognition by CD8^+^ T_M_ cells on dendritic cells initiates their rapid and coordinated production of a burst of CCL3, CCL4 and XCL1 chemokines under the transcriptional control of IRF4. Using intravital microscopy imaging and *in vivo* monoclonal antibody labelling, we reveal that memory CD8^+^ T cells undergo antigen-mediated arrest in splenic red pulp clusters of CCR2^+^ monocytes where they locally deliver both IFNγ- and chemokine-potentiating microbicidal activities to achieve early protection. Thus, rapid and effective memory CD8^+^ T cell responses require a complex series of spatially and temporally coordinated stepwise molecular and cellular events that quickly restrict microbial pathogen growth and optimize the local delivery of effector molecules before clonal expansion occurs.

## Introduction

CD8^+^ T cells have the unique ability to sense and recognize antigens (Ags) derived from intracellular pathogens and tumors (Harty et al., 2000; Vesely et al., 2011; Wong and Pamer, 2003). Live attenuated vaccines using viral backbones (e.g., Vaccinia, Vesicular Stomatitis Virus) or intracellular bacteria such as *Listeria monocytogenes (Lm)* and *Mycobacteria* (BCG), are known to promote robust CD8^+^ T cell responses and establish a pool of systemic and tissue-resident long-lived memory CD8^+^ T (CD8^+^ T_M_) cells. Such CD8^+^ T_M_ cells can rapidly react against immunizing Ags expressed in live vectors, and provide immunity against life-threatening diseases (Martin and Badovinac, 2018; Sallusto et al., 2010; Schenkel and Masopust, 2014).

Cognate Ag, cytokines and chemotactic signals contribute to optimal activation of CD8^+^ T_M_ cells during a recall infection, systemically and at mucosal surfaces (Lauvau et al., 2016; Schenkel and Masopust, 2014). Since MHC class I (MHC-I) molecules are almost ubiquitously expressed, associated Ags can theoretically be presented by any cells, supporting the common view that CD8^+^ T_M_ cells may recognize their cognate Ags and undergo subsequent activation upon triggering by any MHC-I-expressing cells. However, this view was challenged by an important study showing that dendritic cells (DCs) play a unique role in driving optimal Ag-dependent expansion of CD8^+^ T_M_ cells during a recall infection in vaccinated mice (Zammit et al., 2005). The importance of DCs was also extended to the reactivation of tissue-resident memory CD8^+^ T (T_RM_) cells (Shin et al., 2016; Wakim et al., 2008). More recent evidence using two distinct models of lung viral infections, further revealed that draining lymph node (dLN)- derived CD8^+^ T_M_ cells required XCR1^+^ DCs for Ag-dependent reactivation while T_RM_ cells in the lung could be reactivated both by hematopoietic and non-hematopoietic-derived cells (Low et al., 2020). In models of systemic bacterial and viral infections, DCs and Ly6C^+^ CCR2^+^ inflammatory monocytes, are the source of multiple inflammatory cytokines, i.e., IL-12, IL-18, IL-15 and type I interferon (IFN), that drive early Ag-independent, also known as “bystander”, CD8^+^ T_M_ cell-reactivation and differentiation into IFNγ-secreting NKG2D^+^ effector cells (Alexandre et al., 2015; Bedoui et al., 2009; Berg et al., 2003; Raue et al., 2013; Soudja et al., 2012). While such cytokine-driven activation of CD8^+^ T_M_ cells contributes to innate mechanisms of protection, cognate Ag recognition nevertheless remains required to achieve high levels of microbial pathogen-specific immunity. Several mechanisms are likely to account for Ag-dependent CD8^+^ T_M_ cell-mediated protection. These include direct cytolysis of infected cells, secretion of Ag-dependent cytokines (i.e., TNFα) and, very importantly, rapid clonal expansion that ensures sufficient numbers of pathogen-specific effector memory cells are generated to effectively sterilize an infection (Harty et al., 2000; Wong and Pamer, 2003). Trafficking of CD8^+^ T_M_ cells to the sites of infection via chemotaxis (e.g., CXCR3, CCR5) and adhesion (LFA-1, loss of L-selectin), are also essential processes to enable rapid containment and effective elimination of microbial pathogens at portal of entry. Proof of concept studies have used models of systemic viral (Vaccinia, VSV, LCMV) and bacterial (*Lm*) infections in which microbial pathogens are rapidly captured in subcapsular dLNs or splenic marginal zone CD169^+^ macrophages, and drive subsequent homing of CD8^+^ T_M_ cells in response to chemotactic cues (e.g., CXCL9, CXCL10) produced by innate immune and structural cells (Kastenmuller et al., 2013; Maurice et al., 2019; Sung et al., 2012). The massive Ag-independent recruitment of memory cells also leads to inflammation-driven activation of Ag-irrelevant CD8^+^ T_M_ cells (Maurice et al., 2019). While comparable chemotactic mechanisms are also documented in the case of CD8^+^ and CD4^+^ T_RM_ cells in models of skin and vaginal viral infections, initiation of the rapid mucosal immune response by T_RM_ cells is largely dependent on initial cognate Ag recognition, leading to the establishment of a rapid antiviral state that restrict pathogen spreading (Ariotti et al., 2014; Iijima and Iwasaki, 2014; Schenkel et al., 2014; Schenkel et al., 2013). There is, however, still very little knowledge on which early transcriptional gene expression and effector program is specifically triggered in CD8^+^ T_M_ cells upon early cognate Ag recognition and how this enables memory CD8^+^ T cells to mediate the rapid control of pathogen growth and spreading *in situ* in immunized hosts.

Using mice immunized with *Lm*, we previously reported that IFNγ signaling to innate phagocytes, namely CCR2^+^ monocytes and neutrophils, promotes their maturation into TNFα- and reactive oxygen species (ROS)-producing microbicidal effector cells, which accounts for significant protection in both spleen and liver of vaccinated hosts (Narni-Mancinelli et al., 2007; Narni-Mancinelli et al., 2011; Soudja et al., 2014). Yet, in these models of systemic vaccination, as well as others (Kupz et al., 2012; Raue et al., 2013), IFNγ is largely secreted by the CD8^+^ T_M_ cells independent of cognate antigen while protection occurs within hours post challenge infection and before CD8^+^ T_M_ cells even undergo clonal expansion (Narni-Mancinelli et al., 2007; Narni-Mancinelli et al., 2011). In the current work, we dissected the cellular and molecular mechanisms by which cognate antigen programs and orchestrates early CD8^+^ T_M_ cell-mediated pathogen-specific protection in vaccinated hosts undergoing a recall infection. We took advantage of our experimental system in which protection requires cognate Ag recognition but IFNγ, a major protective cytokine *in vivo*, is secreted with no need of cognate Ag, Our results are consistent with a model in which CD8^+^ T_M_ cells migrate to and arrest in infection foci where blood-derived phagocytes, here Ly6C^+^ monocytes, have already been accumulating, to license them with highly effective microbicidal functions for pathogen containment and killing.

## Results

### Cognate antigen *versus* inflammation triggers a broad range of functional pathways in memory CD8^+^ T cells

To understand how cognate antigen orchestrates CD8^+^ T_M_ cell early reactivation and programming *in situ*, we conducted a genome-wide transcriptional analysis of pathogen-specific memory cells undergoing reactivation in the presence or in the absence of their cognate antigen (Ag) (Figure 1). Naïve Ova_257-264_/K^b^-specific OT-I and gB_498-505_/K^b^-specific gBT-I TCR transgenic T cells were adoptively transferred to WT C57BL/6 (B6) mice that were immunized the next day with *Listeria monocytogenes* (*Lm*) expressing both T cell epitopes (*Lm*-Ova-gB). Six weeks later, immunized mice were challenged with *Lm* expressing Ova only (*Lm*-Ova) and we monitored OT-I and gBT-I T_M_ cell activation (Figure 1A). This experimental set-up enabled us to track memory cells that either “see” (OT-I, Ag/Infl.-activated) or do not “see” (gBT-I, Infl.- activated) their cognate Ag. The memory cells were flow-sorted from 8 hr-challenged or control unchallenged mice, and subjected to transcriptomic analysis (Figure 1B). Two-dimensional principal component analysis (PCA) (Figure 1B, left panel) and hierarchical clustering (Figure 1B, right panel) revealed that OT-I T_M_ cells (Ag/Infl.-activated) clustered separately from gBT-I (Infl.-activated) and resting T_M_ (unchallenged) cells that grouped close together. Thus, cognate Ag triggering drives a significantly distinct transcriptional profile in the memory CD8^+^ T cells. A total of 1,837 genes were differentially expressed (p<0.05, fold change>1.5) in activated (Ag/Infl. + Infl.) versus resting T_M_ cells, with the vast majority (1,454, i.e., ∼79%) driven by Ag stimulation only and a smaller proportion triggered by inflammatory signals only (227, i.e., ∼12%) (Figure 1C and Table S1). Only 156 genes (i.e., ∼9%) among the differentially expressed genes, were common between Ag- and inflammation-activated CD8^+^ T_M_ cells. While Ag-stimulation induced similar numbers of up- and down-regulated genes, respectively 703 and 751, inflammation favored the expression of a higher proportion of downregulated genes (152 vs 75 genes) including genes involved in cell adhesion and migration (*Cd44, Cd27, Itgax, S1pr5* ; Figure 1D, Table S1). Commonly genes were similarly distributed between up and downregulation. Further analysis of the genes differentially expressed in Ag- versus inflammation-stimulated CD8^+^ T_M_ cells using biological process gene-ontology (BP-GO) pathway analysis, revealed that cognate Ag, but not inflammation, promoted a wide range of biological functions related to TCR signaling, leukocyte differentiation, apoptosis and cytokine expression (Figure 1E and Table S2). To achieve deeper understanding into the molecular mechanisms by which Ag stimulation modulates the early programming of CD8^+^ T_M_ cells, we plotted the fold change over respective adjusted p-values of all differentially expressed genes (Figure 1F). The most expressed genes in Ag-activated CD8^+^ T_M_ cells encoded for chemokines and cytokines (*Ccl4*, *Ccl3*, *Xcl1* and *Tnfa*), important transcriptional regulators (*Nr4a3*, *Nr4a1*, *Nfat5*, *Zbtb32* and *Irf4*) and proteins involved in proliferation/survival (*Tnfsf14* and *Map2k3*) and cell cycle (*Erg2*). Of note, expression of genes encoding adhesion molecules were largely downregulated (*Itgb6, Itgb3* and *Cdh1*) (Figure 1F). In summary, cognate Ag stimulation endows CD8^+^ T_M_ cells with a robust early multifunctional gene expression program, among which the most significantly upregulated genes encode for chemokines and a specific set of transcription factors.

**Figure 1.**
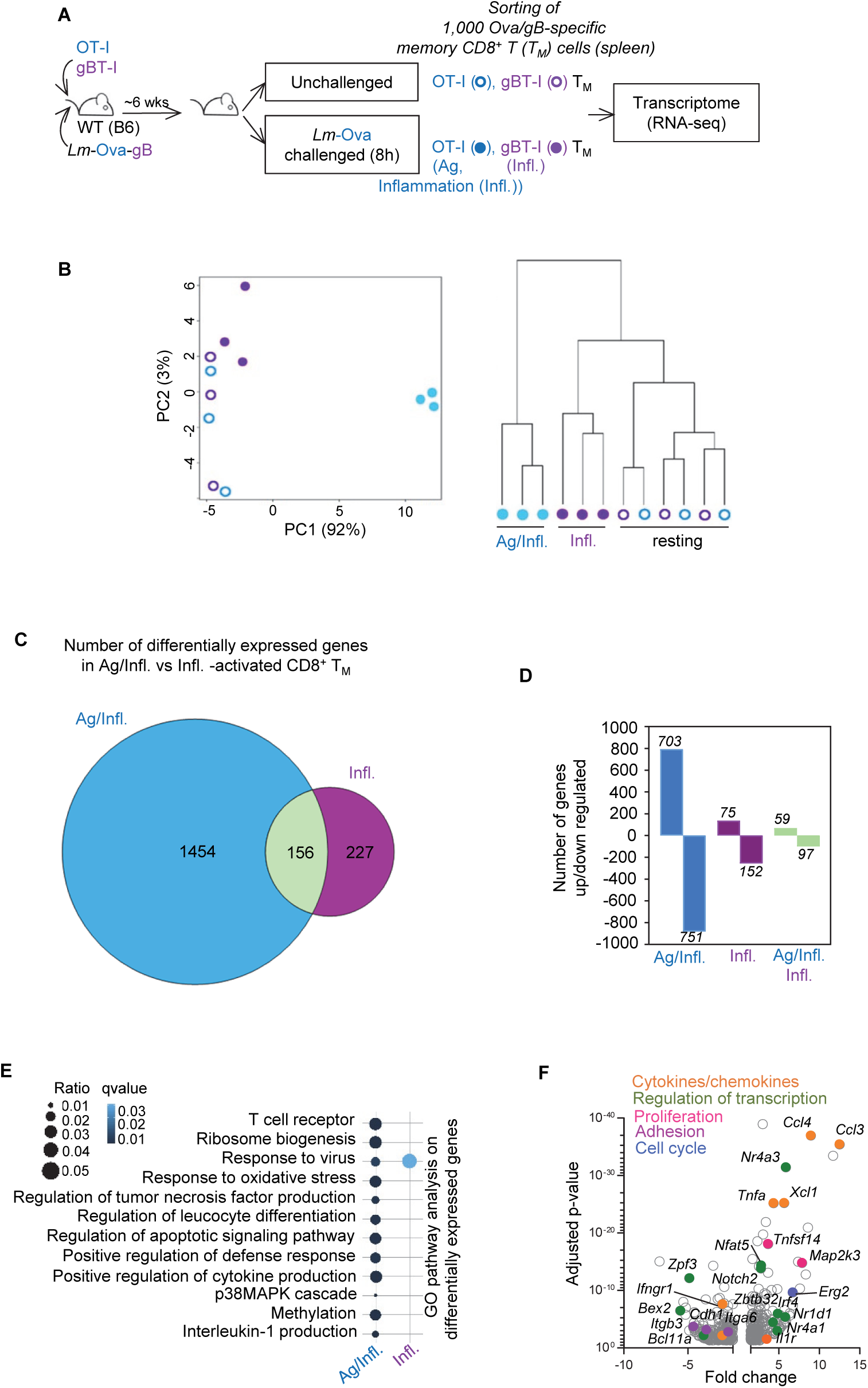
Transcriptomic profiling of antigen/inflammation-*versus* inflammation-activated memory CD8^+^ T cells. (**A**) Schematic of experimental design. Aged-matched WT B6 female mice (CD45.2^+/+^) grafted with tomato-expressing (Td^+^) OT-I and CD45.1^+/+^ gBT-I cells were immunized with 10^4^ *Lm*-Ova-gB, and ∼6 weeks later challenged or not with 10^6^ *Lm*-Ova. After 8 hrs, 1,000 OT-I T_M_ (Ag/Infl, blue) and gBT-I T_M_ (Infl, purple) were flow-sorted from harvested mouse spleens based on CD8, CD3, Tomato (Td^+^) and CD45.1 expression and samples were prepared for RNA-seq analysis. (**B**) PCA plot (left panel) and clustering tree (right panel) of Ag/Infl- (OT-I) *versus* Infl- (gBT-I) stimulated T_M_ cells at steady state and post challenge. Each dot represents an individual mouse and the number in parentheses indicates the percent of variance. Each set of samples (OT-I, gBT-I) was processed in 3 biologically independent replicate experiment. (C) Venn diagrams comparing the numbers of differentially expressed genes Ag/Infl- (OT-I) *versus* Infl- (gBT-I) stimulated T_M_ cells from secondary challenged mice (fold change +/- 1.5, adjusted p-value<0.05). The number of overlapping genes is specified in the green circle. (**D**) Bar graphs representing the number of genes up and down-regulated from the Venn diagram analysis. (**E**) Representation of the top Gene ontology (GO) pathway analysis between Ag/Infl- *versus* Infl-activated T_M_ cells. The size and color of dots are proportional to the number of genes under a specific term and the adjusted p-value, respectively. (F) Volcano plot representing the significantly up and down-regulated genes in Ag/Infl-activated (OT-I) T_M_ cells 8 hrs after the recall challenge infection.

### Memory CD8^+^ T cells produce an early and coordinated burst of chemokines upon cognate antigen recognition

To validate chemokine-encoding gene upregulation in cognate Ag-stimulated CD8^+^ T_M_ cells (Figure 1F), we monitored CCL3, CCL4 and XCL1 chemokine accumulation in Ag (OT-I) *versus* inflammation (gBT-I) triggered T_M_ cells from mice primary immunized with *Lm*-Ova-gB, and challenged 6 weeks later with *Lm-*Ova for 8, 16, 32 and 72 hrs (Figure 2). As early as ∼4 hrs post challenge infection, OT-I, but not gBT-I T_M_ cells, accumulated detectable levels of the 3 chemokines, peaking between 12 and 16 hrs post-infection with 30-40% chemokine^+^ OT-I T_M_ cells (Figure 2A). As expected (Berg et al., 2003; Kupz et al., 2012; Raue et al., 2013; Soudja et al., 2012), both T_M_ cells expressed IFNγ independent of cognate Ag stimulation.Substantial levels of chemokines (CCL3) and IFNγ could be measured in short-term culture supernatants of splenocytes (without brefeldin A) isolated from 8 hrs *Lm*-Ova-challenged *versus* unchallenged mice, indicative of their active secretion (Figure S1A). By 32 hrs, chemokine secretion was terminated, and OT-I T_M_ cells underwent robust clonal expansion (Figure 2B). To further define which subset of CD8^+^ T_M_ cells (Gerlach et al., 2016) among central (CX3CR1^low^CD27^hi^, T_CM_), peripheral (CX3CR1^int^CD27^hi^, T_PM_) or effector (CX3CR1^hi^CD27^low^, T_EM_) CD8^+^ T_M_ cells produced chemokines and IFNγ, we flow-sorted these populations and incubated them with their cognate Ag *in vitro* (Figure 2C and S1B). While both OT-I T_CM_ and T_PM_ accumulated significantly more chemokines and IFNγ than T_EM_ counterparts, they could nevertheless all produce them. To next validate results in endogenous non-TCR transgenic CD8^+^ T_M_ cells and naturally presented epitopes, we immunized WT B6 mice (H2^b^) that also express the K^d^ molecule (B6-K^d^) with *Lm*-gB, allowing for the tracking of both *Lm-*derived LLO_91–99_/K^d^ and p60_217-225_/K^d^ as well as HSV-2-derived gB_497-505_/K^d^ specific CD8^+^ T_M_ cells, using the corresponding tetramers (Tet) (Figure 2D). Six weeks post vaccination, mice were challenged with either *Lm*-gB or *Lm*-LLO_Ser92_ that lacks the LLO_91–99_ epitope, and we monitored endogenous tetramer-specific CD8^+^ T_M_ cell production of chemokines in the presence or the absence of their respective cognate Ags. After *Lm*-gB challenge, e.g., when all T_M_ cell cognate Ags were present, gB_498-505_/K^b^, p60_217-225_/K^d^ and LLO_91-99_/K^d^ tet^+^ CD8^+^ T_M_ cells expressed CCL3. However, when mice were challenged with *Lm*-LLO_Ser92_, inflammation only-stimulated LLO_91-99_/K^d^- and gB_498-505_/K^b^-specific CD8^+^ T_M_ cells expressed IFNγ but no chemokines while Ag-triggered p60_217-225_/K^d^-specific CD8^+^ T_M_ cells accumulated both CCL3 and IFNγ. We next extended findings to CD8^+^ T_M_ cells induced with a different vaccination model, by immunizing mice grafted with OT-I cells with Ova-expressing vesicular stomatitis virus (*VSV-*Ova), challenged them six weeks later with either *Lm*-Ova or *Lm*, and quantified chemokine and IFNγ production (Figure 2E). Likewise upon immunization with *Lm*, CD8^+^ T_M_ cells induced after *VSV* vaccination also induced a rapid and coordinated burst of Ag-dependent chemokines and Ag-independent IFNγ accumulation, peaking at ∼16 hrs post-challenge infection, with 40-60% of chemokine/IFNγ^+^ OT-I T_M_ cells. Thus, altogether these data establish that across distinct mouse models of immunization (bacteria, virus) and multiple CD8^+^ T cell epitopes, cognate Ag recognition triggers a rapid and early coordinated burst of chemokine production by CD8^+^ T_M_ cells.

**Figure 2.**
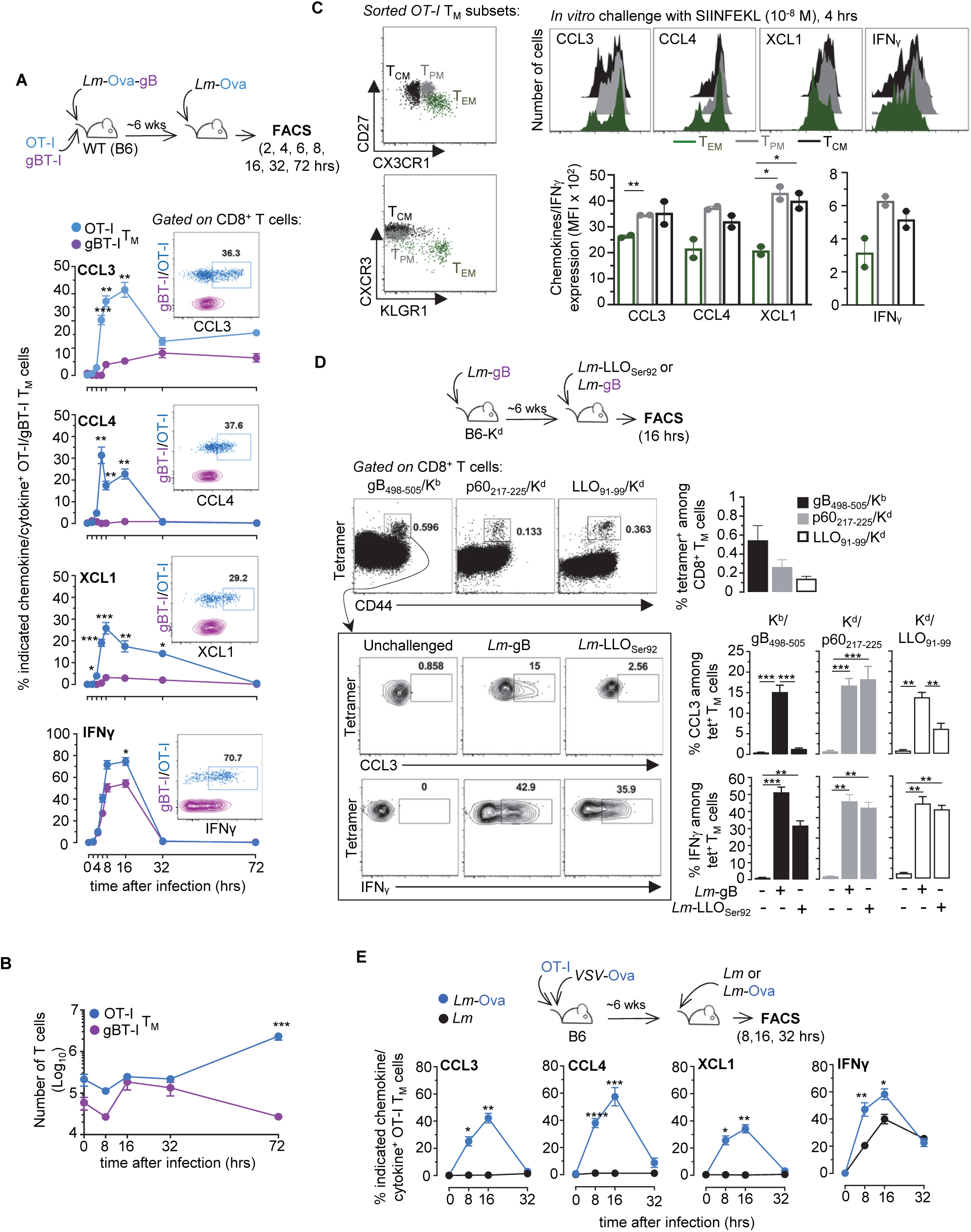
Cognate antigen recognition triggers the rapid and coordinated production of chemokines by memory CD8^+^ T cells. (A-C) WT mice transferred with OT-I Td^+^ and CD45.1^+/+^ gBT-I cells were immunized with 10^4^ *Lm*-Ova-gB and challenged ∼6 weeks later with 10^6^ *Lm*-Ova for 2, 4, 6, 8, 16, 32 and 72 hrs. At indicated times, spleen cell suspensions incubated with Golgi Plug/Stop were stained for cell surface CD8, CD3, CD45.1 and intracellular CCL3, CCL4, XCL1 and IFNγ. (A) Graphs show the kinetic of chemokines and IFNγ accumulation in OT-I (blue) and gBT-I (purple) T_M_ cells and a representative overlaid dot plot of the staining. (B) Number of OT-I (blue) and gBT-I (purple) T_M_ cells at indicated times post recall infection. (C) Subsets of OT-I T_M_ subsets were flow-sorted from the spleens of mice transferred with naïve OT-I cells and immunized with 10^4^ *Lm*-Ova 6 wks before, based on CX3CR1 and CD27 cell surface marker expression (T_EM_: CX3CR1^hi^CD27^low^, T_PM_: CX3CR1^int^CD27^hi^, T_CM_: CX3CR1^low^CD27^hi^). Sorted OT-I T_M_ subsets were next stimulated for 4 hrs with SIINFEKL peptide (10^-8^M) *in vitro* and stained for cell-surface CD8, CXCR3, KLRG1 and intracellular CCL3, CCL4, XCL1 and IFNL. Graphs show expression level of indicated chemokine^+^ and IFN ^+^ OT-I T_M_ subsets with each symbol representing 1 mouse. (D) B6-K^d^ mice were immunized with 10^4^ *Lm*-gB and ∼6 wks later challenged with 10^6^ *Lm*-gB or *Lm*- LLO_Ser92_ for 16 hrs and endogenous memory CD8^+^ T cells were monitored using gB_498-505_/K^b^, p60_217-225_/K^d^ and LLO_91-99_/K^d^ tetramers. Data show the frequency of tetramers^+^ cells among CD8^+^ T_M_ cells and their expression of CCL3 and IFN post challenge with *Lm*-gB or *Lm-* LLO_Ser92_. (E) Mice grafted with OT-I cells were immunized with 2×10^5^ PFU *VSV*-Ova and ∼6 wks later challenged with 10^6^ *Lm* or *Lm*-Ova for 16 hrs. Data show the frequency of CCL3, CCL4, XCL1 and IFNγ among OT-I T_M_ cells after *Lm* or *Lm*-Ova challenge. Representative flow cytometry dot plots are shown. Panels pool data from either 3 independent replicate experiments (A, C) or 2 independent experiments (B, D, E) with n=4-8 mice. P-values (*<0.05), **<0.005, ***<0.0005 and ***<0.0001) are indicated.

### IRF4 exerts transcriptional control over chemokine production by memory CD8^+^ T cells

Cognate Ag stimulation induces upregulation of CCL3, CCL4 and XCL1 chemokine-encoding genes in CD8^+^ T_M_ cells and their subsequent secretion, suggesting a common transcriptional mechanism of regulation. Our transcriptomic analysis revealed several genes involved in the regulation of transcription, such as the transcription factor IRF4, that are upregulated upon cognate Ag recognition. Since IRF4 expression in T cells is directly proportional to the strength of TCR signals (Man et al., 2013; Yao et al., 2013), we expected that, if indeed IRF4 controlled chemokine expression, lowering TCR signaling should lead to a proportional and concomitant loss of IRF4 and chemokine expression by CD8^+^ T_M_ cells. To test this possibility, we used *Lm* expressing three different Ova_257-264_ (SIINFEKL) altered peptide ligands (APLs) in which the original asparagine amino acid in position 4 of the peptide (N4) is replaced by either a glutamine (Q4), a threonine (T4) or a valine (V4), decreasing OT-I TCR signaling by factors of ∼20, 70 and 700 times, respectively (Zehn et al., 2009). Mice grafted with OT-I and gBT-I cells were immunized with *Lm*-Ova-gB, and 6 weeks later, either left unchallenged or challenged with *Lm* expressing each Ova APL or control *Lm*-Ova (N4). We next monitored the secretion of chemokines and IFNγ 16 hrs later (Figure 3A). Decreasing OT-I TCR signaling led to a proportional loss of chemokine-producing T cells (CCL3, CCL4 and XCL1), which also directly correlated with the loss of IRF4 expression (Figure 3B). As expected, inflammation-stimulated gBT-I T_M_ cells neither produced chemokines nor expressed IRF4, while IFNγ production remained comparable across all challenge conditions, in both cognate Ag (OT-I) and inflammation (gBT-I) triggered CD8^+^ T_M_ cells. To ensure that IRF4 up-regulation during endogenous pathogen-specific polyclonal CD8^+^ T_M_ cell response, was comparable to that of OT-I TCR transgenic T cells, we next used the same immunization/challenge approach as in Figure 2C. Here, gB_498-505_/K^b^, p60_217-225_/K^d^ and LLO_91-99_/K^d^ tet^+^ CD8^+^ T_M_ cells underwent the most robust upregulation of IRF4 expression during challenge infection in presence their respective cognate Ag (Figure 3C). These results collectively indicate a direct correlation between the strength of TCR signaling and the proportion of chemokine-producing CD8^+^ T_M_ cells. Furthermore, *in vitro* “challenge” of OT-I T_M_ cells isolated from *Lm-*Ova-immunized mice with the SIINFKEL epitope in the presence or absence of broad inhibitors of either translation (cycloheximide) or transcription (Actinomycin D), suggested that most of the CCL3 in Ag- stimulated T_M_ cells was being rapidly transcribed (>60%)(Figure 3D) while only a smaller proportion was stored as mRNA (∼30%) but none as protein, a result also consistent with recent reports (Davenport et al., 2020; Eberlein et al., 2020). Taken together, these data support the hypothesis that IRF4 acts as a transcriptional regulator of chemokine expression downstream of TCR signaling.

**Figure 3.**
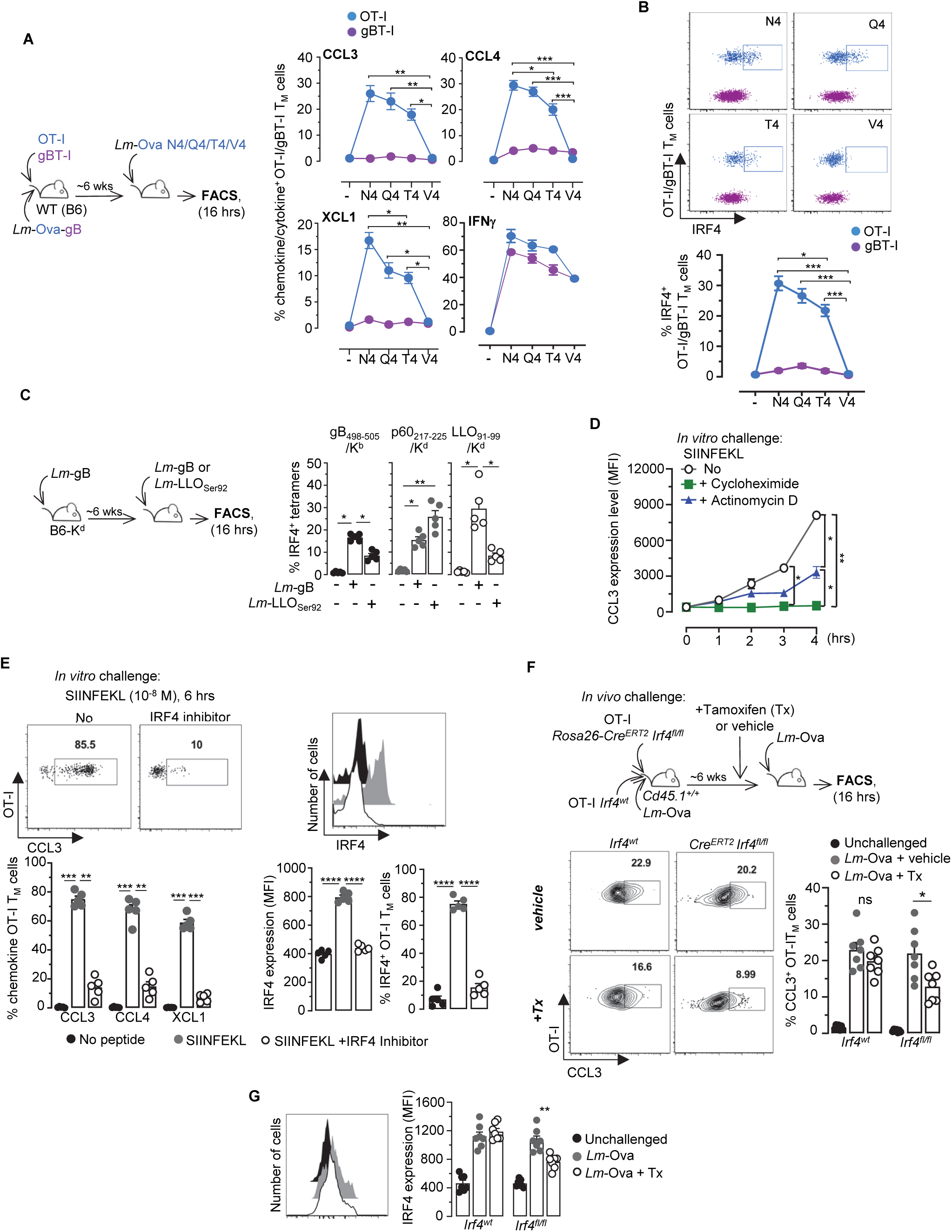
The transcription factor IRF4 orchestrates chemokine production by memory CD8^+^ T cells downstream TCR-signaling. (A-B) Mice grafted with OT-I Td^+^ and CD45.1^+/+^ gBT-I cells were immunized with 10^4^ *Lm*-Ova-gB, and ∼6 wks later challenged or not for 16 hrs with 10^6^ *Lm*-Ova N4 or *Lm* expressing 3 different Ova APLs, namely *Lm-*Ova Q4, *Lm-*Ova T4 or *Lm-*Ova V4. Spleen cell suspensions were next incubated with Golgi Plug/Stop and stained for cell-surface CD8, CD3, CD45.1 and indicated intracellular chemokines and IFNγ (A), or IRF4 (B). Graphs represent the proportion of OT-I or gBT-I T_M_ cells expressing indicated intracellular markers after challenge with *Lm*-expressing N4, Q4, T4 or V4. Representative overlaid dot plots of IRF4 intracellular staining in OT-I and gBT-I T_M_ cells are shown in (B) . (C) B6-K^d^ mice were immunized with 10^4^ *Lm*-gB and ∼6 weeks later challenged with 10^6^ *Lm*-gB or *Lm*-LLO_Ser92_ for 16 hrs, and endogenous memory CD8^+^ T cells were quantified using gB_498-505_/K^b^, p60_217- 225_/K^d^ and LLO_91-99_/K^d^ tetramers. Bar graphs indicate the proportion of IRF4^+^ cells among CD8^+^ tetramer^+^ cells. (D, E) Splenocytes from 6 wks-immunized mice following the experimental design depicted in (A), were incubated for 6 hrs with SIINFEKL (10^-8^M) *in vitro* with or without either cycloheximide or actinomycin D for indicated times (D), or with an IRF4 inhibitor (SCG-CBP30, 20µM) for 6 hrs (E). OT-I Td^+^ T_M_ cells were stained for cell-surface CD8, CD3 and intracellular CCL3, CCL4 and XCL1 and IRF4. Graphs show the proportions and/or expression level of indicated chemokine^+^ and IRF4^+^ OT-I T_M_ cells. (F, G) *Rosa26^CreERT2^Irf4^flox/flox^Cd45.2^+/+^* and WT *Cd45.1^+/-^* OT-I cells were co-transferred to *Cd45.1^+/+^* WT recipient mice and immunized with 10^4^ *Lm*-Ova the next day. Six wks later, mice received Tx (1mg/day) or vehicle i.p. every day for 5 days before secondary challenge infection with 10^6^ *Lm*-Ova. At 16 hrs, CCL3 and IRF4 expression was determined as described above in *Irf4^flox/flox^ versus* WT OT-I T_M_ cells in the different experimental conditions. Representative FACS dot plots and histograms staining are shown. Panels pool the result of 2 independent replicate experiments with n=6 (A, B), 5 (C-E) and 7 (F,G) mice. P-values are indicated.

To establish whether IRF4 controls CCL3, CCL4 and XCL1 chemokine expression in CD8^+^ T_M_ cells, we blocked IRF4 in OT-I T_M_ cells, and quantified their production of chemokines. Through *in vitro* SIINFKEL peptide challenge of OT-I T_M_ cells isolated from *Lm-*Ova-immunized mice with the chemical inhibitor SCG-CBP30, which selectively inhibits bromodomain-containing transcription factors like IRF4 (Figure 3E), we found that IRF4 expression in OT-I T_M_ cells was prevented, and the proportion of chemokine^+^ cells was significantly decreased (by ∼80%) compared to incubation with peptide-only. To confirm and validate findings *in vivo*, we generated OT-I^+^ *Rosa26^CreERT2^Irf4^flox/flox^* mice in which *Irf4* could be inducibly deleted in OT-I T_M_ cells (Figure 3F). Naïve *Rosa26^CreERT2^Irf4^flox/flox^* or WT *Cd45.1/2* OT-I cells were co-transferred to WT *Cd45.1^+/+^* recipient mice then immunized with *Lm*-Ova, and 6 weeks later, mice either received tamoxifen (Tx) or vehicle for 5 days before *Lm*-Ova recall infection. In Tx-treated groups, the proportion of *Rosa26^CreERT2^Irf4^flox/flox^* OT-I T_M_ cells secreting chemokines (CCL3), was significantly decreased compared to that of WT counterparts (by ∼45%), yet both of these genotypes secreted comparable amounts of CCL3 in mock-treated mice. Expression levels of IRF4 was also diminished in Tx- but not mock-treated *Rosa26^CreERT2^Irf4^flox/flox^* OT-I T_M_ cells, further validating this result (Figure 3G). In conclusion, our data demonstrate that IRF4 is a master transcriptional regulator of the coordinated and simultaneous burst of CCL3, CCL4 and XCL1 chemokines produced by Ag-activated CD8^+^ T_M_ cells *in vitro* and *in vivo*.

### Monocyte clustering occurs independent from cognate antigen or IFNγ-signaling

We previously showed that CD8^+^ T_M_ cell-mediated control of *Lm* growth during recall infection occurs within only few hours (∼6-8 hrs) and correlates with their rapid localization with clustered Ly6C^+^/CCR2^+^ monocytes and neutrophils in the splenic red pulp (RP) of infected mice, at portal of bacterial entry (Bajenoff et al., 2010; Narni-Mancinelli et al., 2011; Soudja et al., 2014). Thus, we hypothesized that CD8^+^ T_M_ cell-derived chemokines produced in response to cognate Ag recognition, orchestrate monocyte homing and clustering to rapidly prevent pathogen spreading and help deliver local IFNγ. To gain deeper understanding of this process, we first monitored the kinetic of Ly6C^+^ monocyte clustering in the RP of *Lm-*vaccinated mice undergoing a recall infection (Figure S2A). *Ccr2*^CFP^ mice, in which all CCR2^+^Ly6C^+^ monocytes express the CFP reporter protein, were grafted with OT-I cells and immunized with *Lm-*Ova. Six weeks later, mice were left unchallenged or challenged with *Lm*-Ova and spleens harvested 3, 6, 16 and 40 hrs later for whole organ tile reconstruction using multi-photon laser scanning microscopy that only enables to visualize splenic RP (Figure S2A). Already by 3 hrs post-challenge infection, few clusters of CCR2^+^ monocytes were detected, with their proportion increasing from 6 hrs to their peak at 16 hrs, and with clusters still present by 40 hrs. Of note, peak clustering of monocytes at 16 hrs correlated with that of chemokines produced by the T_M_ cells (Figures S2A and 2A). Unexpectedly, however, unimmunized mice challenged with *Lm*- Ova, which do not control the infection compared to immunized counterpart (Figure S2B), still developed comparable numbers and volume of CCR2^+^ monocyte clusters 16 hrs post-infection (Figure 4A). This result indicated that the presence of immunization-induced memory cells is not essential for monocyte homing and clustering to occur, though they may still alter monocyte functions. To better investigate the role of CD8^+^ T_M_ cells and cognate Ag in monocyte cluster formation, we adoptively transferred OT-I T_M_ cells in *Ccr2*^CFP^ WT mice subsequently challenged with *Lm* or *Lm*-Ova, and monitored monocyte clustering in spleen RP (Figure 4B). For this, we took advantage of a heterologous prime/boost immunization strategy of mice grafted with OT-I cells, primed with *VSV*-Ova, and challenged with *Lm*-Ova to generate sufficiently high numbers of OT-I T_M_ cells for purification and transfer. Whether cognate Ag was present or not, the proportion and volume of monocyte clusters at the peak (16 hr) remained comparable, a result that we also confirmed in WT *Ccr2*^CFP^ mice grafted with OT-I cells, immunized with *VSV*-Ova and challenged with either *Lm* or *Lm*-Ova 6 weeks later (Figure S3A). Lastly, since IFNγ signaling is an essential contributor to vaccinated host protective responses (Soudja et al., 2014), we tested if it may direct monocyte clustering. For this, we adoptively transferred OT-I T_M_ cell in *Ifngr1^-/-^* mice that we next challenged with *Lm* or *Lm*-Ova (Figure 4C). Since monocytes in *Ifngr1^-/-^* mice did not express CFP, we tracked them using intravenous injection of anti-Ly6C-PE mAb 16 hrs prior to imaging, which co-labels all detectable clustered CFP^+^ monocytes (Figure S3B). As before, whether cognate Ag (Ova) and IFNγ signaling were present or not, the proportion and volume of monocyte clusters at 16 hrs were also comparable. Hence, taken together, these data establish that monocyte homing and clustering occurs largely independent of the presence of Ag-specific CD8^+^ T_M_ cells and IFNγ signaling.

**Figure 4.**
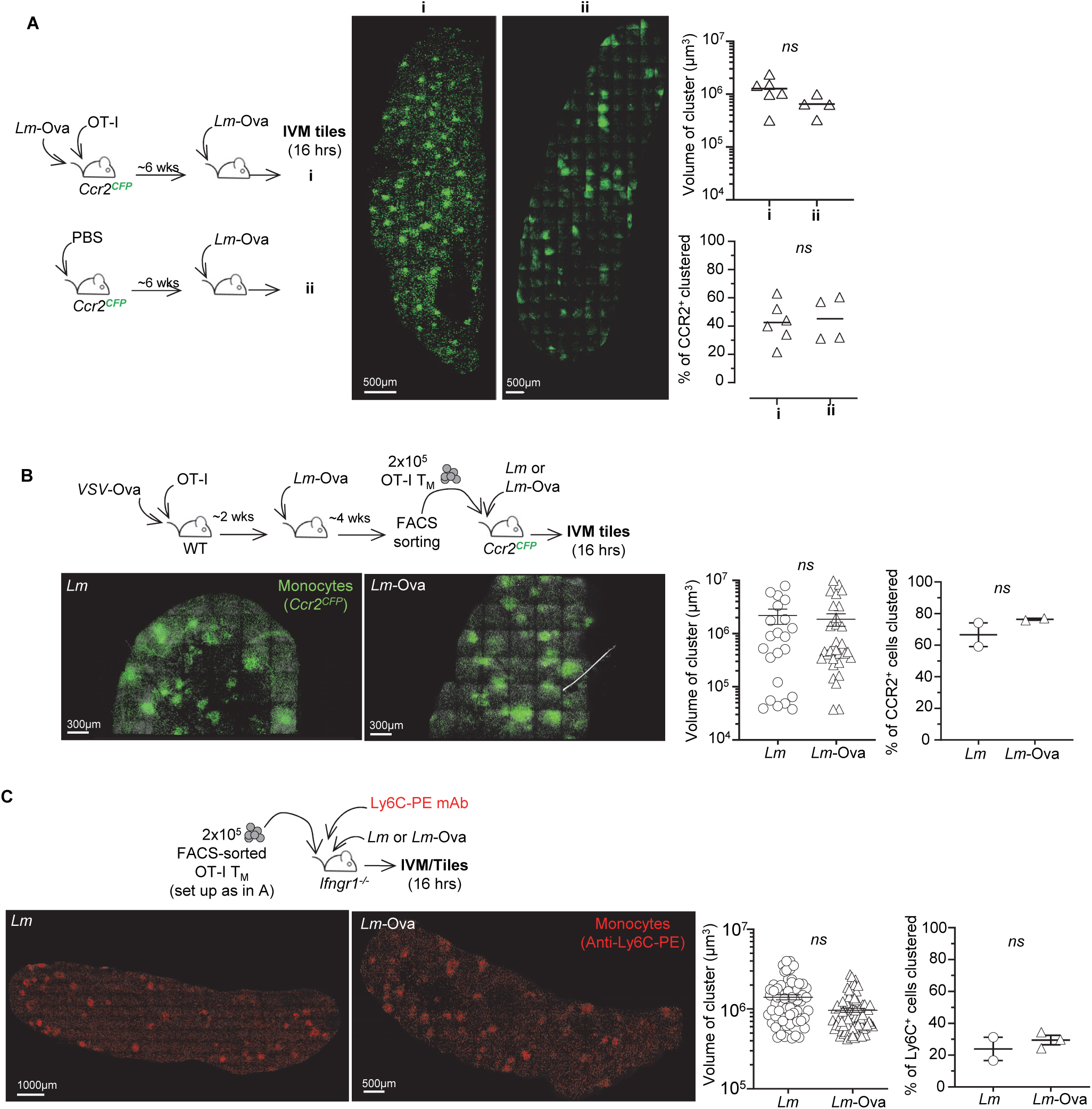
Ly6C/CCR2^+^ monocyte clusters form in the splenic red pulp independent from memory CD8^+^ T cells and IFNγ. (A) Representatives IVM tiles of reconstructed *Ccr2^CFP^* mouse spleens. As depicted in the schematics, mice were transferred with OT-I cells and primary and secondary challenged with *Lm*-Ova (panel i) or only primary immunized with *Lm*-Ova. Ccr2^CFP^ monocytes are in green and scales are indicated. Graphs show the average volume of clusters and proportion of clustered monocytes in each mouse spleen analyzed. (B, C) 2×10^5^ OT- I T_M_ flow-sorted cells generated upon immunization with 2×10^5^ PFU *VSV*-Ova and challenge with 10^6^ *Lm*-Ova, were transferred to naive *Ccr2^CFP^* (B) or *Ifngr1^-/-^* (C) recipient mice subsequently challenged with 10^6^ *Lm* or *Lm*-Ova for 16 hrs. Representative IVM tiles of reconstructed mouse spleens with CCR2^+^ (B) or Ly6C^+^ (C) monocytes in spleen’s red pulp. In (C) *Ifngr1^-/-^* were also co-injected with anti-Ly6C-PE ab (10µg). Graphs in (B, C) show the volume of individual clusters and the average proportion of clustered Ly6C^+^ monocytes in each mouse spleen analyzed across 2 independent replicate experiments (n=2-3).

### Cognate antigen on dendritic cells but not monocytes, control CD8^+^ T_M_ cell-production of chemokines and arrest in monocyte clusters

While monocyte homing and clustering still occurred in unimmunized mice, these clusters could nevertheless be necessary to mount a protective recall response in immunized mice. Thus we pursued the hypothesis that monocyte clustering is functionally important, and may act as local “hubs” in which CD8^+^ T_M_ cells arrest and deliver IFNγ and other effector molecules to them, as well as to other innate immune cells recruited to these clusters -i.e., neutrophils, NK cells (Bajenoff et al., 2010; Soudja et al., 2014). We used intravital imaging microscopy (IVM) of spleen RP in *Ccr2*^CFP^ living mice undergoing a recall infection (Figure 5A and Movies S1, S2, S3). Mice transferred with OT-I (Td^+^) and gBT-I (GFP^+^) cells, were immunized with *Lm-*Ova-gB and challenged 6 weeks later with either *Lm*-Ova, *Lm* or *Lm*-Ova-gB. In *Lm*-Ova-challenged mice, in which only OT-I T_M_ cells recognize their cognate Ag, most OT-I T_M_ cells localized in the cluster of monocytes (CFP^+^) and arrested, or only exhibited very limited motility (Track velocity 1.93 µm/min) (Figures 5A, Movie S1 and Figure S4A). In contrast, gBT-I T_M_ cells were more motile (track velocity 4.01 µm/min), yet were enriched in the monocyte clusters similarly to OT-I cell counterparts (Figure S4B). Both T_M_ cells speed also decreased inside compared to outside monocyte clusters, collectively suggesting that non-cognate Ag signals impact their homing to and motility in the clusters (Figure 5A). As expected, in *Lm*-Ova-gB-challenged mice, where both T_M_ cells recognize their cognate Ag, OT-I and gBT-I T_M_ cells arrested in the clusters while simultaneously exhibiting higher motility outside of clusters (Movie S2 and Figure S4B). Moreover, in *Lm*-challenged mice in which no cognate Ag was present, both T_M_ cells exhibited the same pattern of enriched localization inside versus outside the clusters, and comparable speeds (Movie S3 and Figure S4A, B). Thus, cognate-Ag signals induce Ag-specific T_M_ cell arrest in Ly6C/CCR2^+^ monocyte clusters where IFNγ is detected in T_M_ cells (Bajenoff et al., 2010; Soudja et al., 2014), indicating that a potential functional interaction between T_M_ cells and monocytes is occurring in these clusters. In addition, the fact that even non-cognate Ag-specific T_M_ cell speed was reduced inside compared to outside of clusters, suggested that the clusters were conductive of a qualitatively distinct, possibly hypoxic, local microenvironment (Waite et al., 2011).

**Figure 5.**
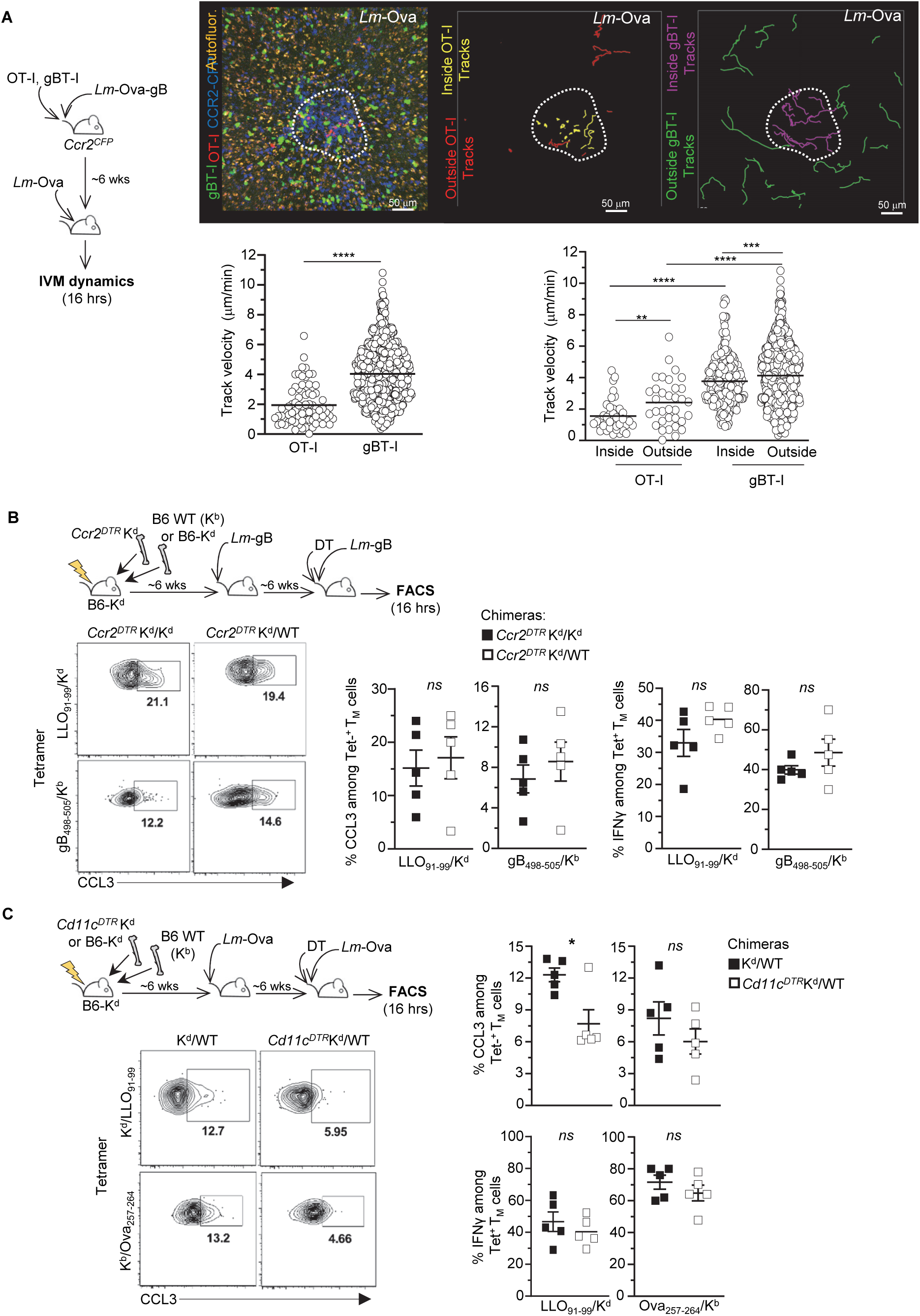
Memory CD8^+^ T cells arrest upon cognate Ag recognition presented by dendritic cells by not CCR2^+^ monocytes. (A) *Ccr2^CFP^* mice co-transferred with naïve OT-I Td^+^ and gBT- I GFP^+^ cells and immunized with 10^4^ *Lm-*Ova-gB were challenged ∼6 wks later with *Lm*-Ova, and spleens from live mice were exposed and imaged 16 hrs later using IVM imaging. A representative image (left) of OT-I (red) and gBT-I (green) T_M_ cell localization in a cluster (white dotted line) of CCR2^+^ monocytes (blue) is shown. Autofluorescence appears in yellow. Also shown (center and right images) are OT-I T_M_ cell tracks (outside, red and inside, yellow) and gBT-I T_M_ cell tracks (outside, green and inside, purple) inside/outside the same cluster of CCR2^CFP^ monocytes. Graphs represent the speed of individual OT-I and gBT-I T_M_ cells in the monocyte cluster area (left) and inside/ outside the cluster. (B, C) Lethally irradiated (1,200 rads) B6-K^d^ recipient mice were reconstituted with (B) B6-K^d^ or WT B6 (K^b^) and *Ccr2^DTR^* K^d^ BM or (C) *Cd11c^DTR^*K^d^ or K^d^ and WT B6 BM. Six weeks post reconstitution, mice were immunized with 10^4^ *Lm-*gB (B) or *Lm*-Ova (C) and 6 weeks later, challenged with 10^6^ *Lm-*gB 12 hrs post diphtheria toxin (DT)-treatment. Endogenous CD8^+^ T_M_ cells were monitored using LLO_91-99_/K^d^, gB_498-505_/K^b^ or Ova_257-264_/K^b^ tetramers. Graphs show the expression of CCL3 and IFN among tetramers^+^ (Tet^+^) cells after challenge. Each symbol corresponds to 1 individual mouse in 1 of 2 replicate experiment and p-value are shown.

Since T_M_ cells arrest in monocyte clusters in the presence of cognate Ag, and T cells arrest in response to Ag recognition (Bousso and Robey, 2003; Mempel et al., 2004; Stoll et al., 2002), we postulated that monocytes may present Ag to them. To test this possibility, we generated mixed bone-marrow (BM) chimera mice in which selective elimination of K^d^-dependent cognate Ag presentation by Ly6C^+^ monocytes can be achieved. Here, lethally irradiated K^d^-expressing B6 mice (B6-K^d^) were reconstituted with *Ccr2^DTR^* K^d^ BM and either i) B6-K^d^ (K^d^) or ii) B6 (WT) BM (1:1 ratio), producing *Ccr2^DTR^* K^d^/WT mice and *Ccr2^DTR^* K^d^/K^d^ chimeras. In these mice, diphtheria toxin (DT) injection eliminates CCR2^+^K^d^ monocytes while DTR^neg^ (K^d^ or WT) CCR2^+^ monocytes remain, respectively (Figure S3C). Chimeras were immunized with *Lm*-gB and treated with DT prior to *Lm*-gB challenge infection, and we monitored both LLO_91-99_/K^d^ and gB_498-505_/K^b^ Tet^+^ CD8^+^ T_M_ cells for Ag-dependent chemokine (CCL3) and Ag-independent IFNγ production (Figure 5B). The proportion of Ag-stimulated (CCL3^+^) LLO_91-99_/K^d^ tet^+^ CD8^+^ T_M_ cells was the same whether CCR2^+^ monocytes could present the LLO_91-99_/K^d^ Ag (in DT-treated *Ccr2^DTR^* K^d^/K^d^ chimeras) or not (in DT-treated *Ccr2^DTR^* K^d^/WT chimeras). Yet, the frequency of IFNγ^+^ cells was equivalent, confirming that LLO_91-99_/K^d^ tet^+^ CD8^+^ T_M_ cells underwent comparable Ag-independent activation in all groups. No differences in the proportion of CCL3^+^ and of IFNγ^+^ gB_498-505_/K^b^ tet^+^ CD8^+^ T_M_ cells were measured between the various experimental conditions, ruling out a possible impact of DT-induced deletion on T_M_ cell activation. Thus, Ag presentation by splenic CCR2^+^/Ly6C^+^ monocytes is not required for Ag-dependent CD8^+^ T_M_ cell-activation during recall infection.

Dendritic cells (DCs) quickly uptake *Lm* (Edelson et al., 2011; Neuenhahn et al., 2006) and contribute to CD8^+^ T_M_ cell-reactivation (Zammit et al., 2005). Using K^d^/WT and *Cd11c^DTR^* K^d^/WT chimera mice, in which DT injection eliminates CD11c^+^K^d^ DCs while DTR^neg^ (WT or K^d^) CD11c^+^ DCs remain (Figure S3D), we tested whether CD11c^+^ DC presented cognate Ag to T_M_ cells after immunization/challenge with *Lm*-Ova (Figure 5C). A significant decrease (∼40%) in CCL3^+^ CD8^+^ T_M_ cells was only measured for LLO_91-99_/K^d^ but not Ova_257-264_/K^b^ tet^+^ CD8^+^ T_M_ cells, while the proportion of IFNγ^+^ cells remained equivalent between the different groups of chimeras. Thus, taken together, these data indicate that splenic CD11c^+^ DCs but not CCR2^+^ monocytes, selectively present cognate Ag to CD8^+^ T_M_ cells.

### Cognate Ag stimulation of CD8^+^ T_M_ cells potentiates monocyte effector functions in the clusters

Cognate Ag enables CD8^+^ T_M_ cell arrest in monocyte clusters and their concomitant production of a chemokine burst. If, as hypothesized, CD8^+^ T_M_ cell arrest in these clusters is functionally important for local delivery of chemokines and IFNγ, we predicted that in the presence of cognate Ag, these cells should produce more effector cytokines (Figure 6A). To test this model, we immunized mice transferred with OT-I T_M_ cells with *VSV*-Ova. Six weeks later, mice were challenged with either *Lm*-Ova or *Lm*, and we monitored TNFα and CXCL9 production in monocytes and neutrophils. In the presence of cognate Ag recognition (*Lm*-Ova challenge), the proportion of monocytes and neutrophils producing TNFα and CXCL9 was significantly increased (factor of ∼3) compared to mice challenged without cognate Ag (*Lm*). Interestingly, CCR5 and XCR1 expression on monocytes (but not on neutrophils) was significantly increased after the challenge infection, suggesting monocytes become more responsive to chemokine signaling during infection (Figure 6B). We next directly assessed if adding recombinant CCL4 or XCL1 chemokines, which respectively bind CCR5/CCR1 and XCR1, to monocytes and neutrophils isolated from secondary challenged mice, enhanced their production of TNFα *ex vivo*. After incubation with rCCL3, rCCL4 and rXCL1, monocytes from challenged mice accumulated intracellular TNFα in 15, 20 and 40% of total monocytes, respectively, and in a dose-dependent manner (Figure 6C and S5A, B). Incubation of naïve splenocytes with rCCL3 also induced a significant proportion of TNFα^+^ monocytes (∼30%) compared to unstimulated counterparts, though this was substantially less than after heat-killed *Lm* (HK*Lm*) stimulation, a known robust trigger of monocyte activation (Figure S5C). Neutrophils, however, largely failed to respond to chemokine restimulation *ex vivo*, in line with their low levels of cell-surface expression of chemokine receptors (10-15 times lower than monocytes), rather implicating a chemokine-independent mechanism for their activation (Figure S5D). Blocking CCR5 and CCR1 with chemical inhibitors during co-incubation with recombinant chemokines prevented TNFα production by monocytes, ruling out any CCR1/CCR5-independent activation mechanisms (Figures 6C and S5A). Incubation with HK*Lm* induced 40-50% of them to express TNFα, a proportion similar to that measured in monocytes incubated with rXCL1 or the combination of chemokines.

**Figure 6.**
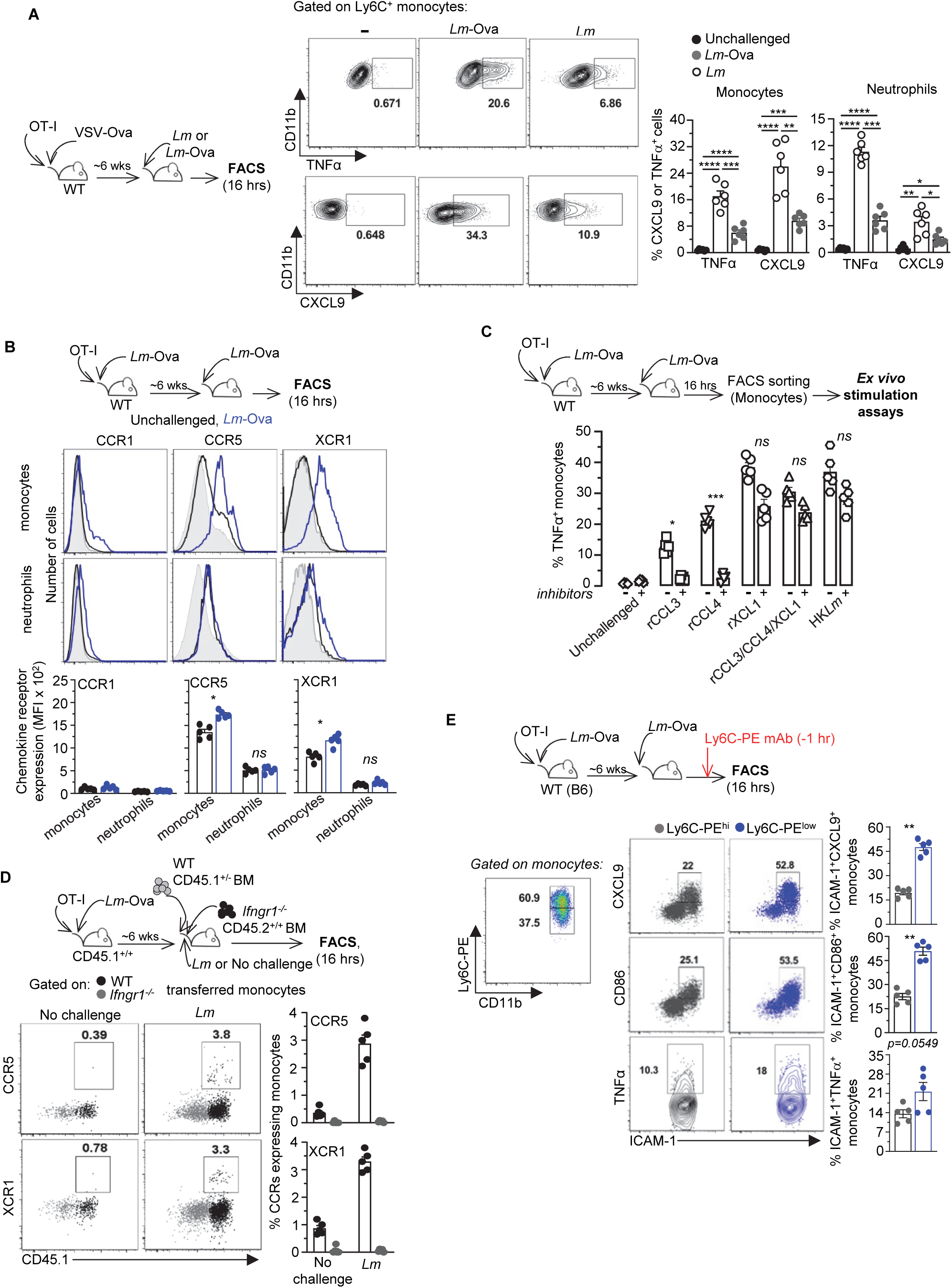
Cognate Ag and chemokine signaling enhance monocyte effector functions. WT mice transferred with OT-I cells were immunized with *VSV-*Ova (A) or *Lm-*Ova (B, C) and challenged or not, 6 wks later with *Lm* or *Lm-*Ova. Spleens from 16 hr-challenged or unchallenged mice were harvested, and cells incubated for 4-6 hrs with brefeldin A prior to staining for expression of CD11b, Ly6C and Ly6G cell surface markers and (A) indicated intracellular effector and chemotactic markers or (B) expression of CCR1, CCR5 and XCR1 chemotactic receptors. In (C), monocytes and neutrophils from the spleens of challenged mice were flow-sorted and either stimulated or not with indicated recombinant chemokines or HK*Lm*, with or without CCR5 and CCR1 inhibitors, prior to staining for intracellular TNFα. (D) BM from WT (CD45.1^+/-^) and *Ifngr1^-/-^* (CD45.2^+/+^) mice were co-transferred to CD45.1^+/+^ recipient mice immunized with *Lm*-Ova 6 weeks before, and immediately challenged with *Lm*. 16 hrs later, spleen cells were stained for expression of CCR5 and XCR1 on monocytes. (E) *Lm-*Ova immunized mice were challenged with *Lm*-Ova or not, and 1 hr before sacrifice, 5 μg Ly6C-PE mAb was injected i.v. to the hosts. Spleens were harvested and cells stained for cell surface CD11b, Ly6C-PerCpCy5.5, ICAM-1, CD86 and intracellular TNFα and CXCL9. After gating on Ly6C-PerCpCy5.5^+^ monocytes, Ly6C-PE^hi^ and Ly6C-PE^low^ monocytes were identified and further analyzed for indicated marker expression. Representative FACS dot plots are shown and bar graphs pool 2 independent replicate experiments with n=6 (A, D) and 5 (B, C and E) mice. P- values are indicated.

Since both cognate Ag and CD8^+^ T_M_ cell-derived IFNγ are required for optimal monocyte production of TNFα (Figure 6A, (Soudja et al., 2014)), we further hypothesized that IFNγ signaling triggers chemokine receptor upregulation on monocytes, making them more responsive to the chemokines released upon cognate Ag stimulation (Figure 6D). We tested this idea by co-transferring WT and *Ifngr1^-/-^* BM into WT recipient mice immediately challenged with *Lm*, and monitored CCR5 and XCR1 chemokine receptor expression 16 hrs later. While we could detect CCR5- and XCR1-expressing WT monocytes, *Ifngr1^-/-^* monocytes failed to upregulate expression of these receptors, indicating that IFNγ signaling likely modulates cell-surface expression of CCR5 and XCR1 on monocytes. Hence, these results collectively establish that the same chemokines produced by Ag-stimulated CD8^+^ T_M_ cells can directly signal to chemokine receptors expressed on monocytes, to further enhance their ability to produce TNFα, a cytokine that is absolutely required for host protective memory responses against secondary *Lm* infection (Narni-Mancinelli et al., 2007; Neighbors et al., 2001).

We next formally asked whether Ly6C^+^ monocyte activation *in vivo* was spatially restricted to their clusters. For this, we sought to measure the activation of monocytes inside versus outside the clusters. Similar *Lm*-induced clusters of myeloid cells have been reported to exclude dextrans suggesting they were not diffusive (Waite et al., 2013). Therefore, we stained monocytes *in vivo* using anti-Ly6C-PE mAb injected 1 hr prior to spleen harvest (1 hr labelling), which we found labels all Ly6C^+^ splenocytes that are not within established clusters, in contrast to injecting anti-Ly6C-PE mAb at the time of challenge infection (16 hrs labelling), prior to cluster formation (Figures S6A and S3B). With the 1 hr labelling approach, >90% of Ly6C^+^ monocytes exhibited equivalent Ly6C-PE staining in unchallenged mice (no clusters), while ∼40% of them had lower Ly6C-PE staining in challenged mice, a proportion consistent with that of clustered monocytes in our microscopy quantifications (Figures S6B, 4A and S2A). With this approach, we could determine whether monocyte activation was dependent on localization within clusters during recall infection (Figure 6E and S6C). A significantly higher proportion of Ly6C-PE^low^ (clustered) compared to Ly6C-PE^hi^ (non-clustered) monocytes expressed higher levels of ICAM-1, CD86 and intracellular CXCL9 and TNFα, demonstrating that monocytes undergo robust activation within the clusters, and consistent with local delivery of activating chemokines and IFNγ by CD8^+^ T_M_ cells.

### Cognate antigen stimulation and IFN**γ** signaling are both required for memory CD8^+^ T cell-dependent protection of immunized mice

Since both cognate Ag stimulation and IFNγ-signaling are required for CD8^+^ T_M_ cell-dependent protection of immunized hosts against challenge infection, we next assessed the relative contribution of both mechanisms. We adoptively transferred OT-I T_M_ cells to naïve WT or *Ifngr1^-/-^* mice that were further challenged with a lethal dose of *Lm* (no cognate Ag) or *Lm*- Ova (with cognate Ag)(Figure 7A). Control groups did not receive any OT-I T_M_ cells. Bacterial titers in spleens and livers were next quantified 24 hrs later. While as expected, transfer of OT-I T_M_ cells confer significant levels of protection to WT recipient mice against *Lm*-Ova challenge (considered 100%), protection was reduced to ∼40% in both organs when challenged with *Lm* (no cognate Ag). These findings were also recapitulated in WT mice primary immunized with *VSV*-Ova, and challenged 6 weeks later with either *Lm* or *Lm*-Ova (Figure S5A). Interestingly, however, OT-I T_M_ cell-transfer in *Ifngr1^-/-^* mice only conferred modest protection against challenge with *Lm* or *Lm*-Ova, with over 60% protection loss compared to WT mice. Yet, in *Lm-* Ova (but not *Lm*) challenged *Ifngr1^-/-^* mice, OT-I T_M_ cells still efficiently recognized their cognate Ag and produced chemokines (Figure S7B). Consistent with these results, monocyte and neutrophil production of TNFα and CXCL9 effector cytokine/chemokine in WT or *Ifngr1^-/-^* mice that received OT-I T_M_ cells was significantly reduced when cognate Ag was absent (*Lm* challenge) or IFNγ-signaling (*Ifngr1^-/-^*) was disrupted (Figure 7B). Taken together, these results indicate that cognate Ag stimulation and IFNγ-signaling are both required to achieve optimal protection and that neither of these signals are individually sufficient.

**Figure 7.**
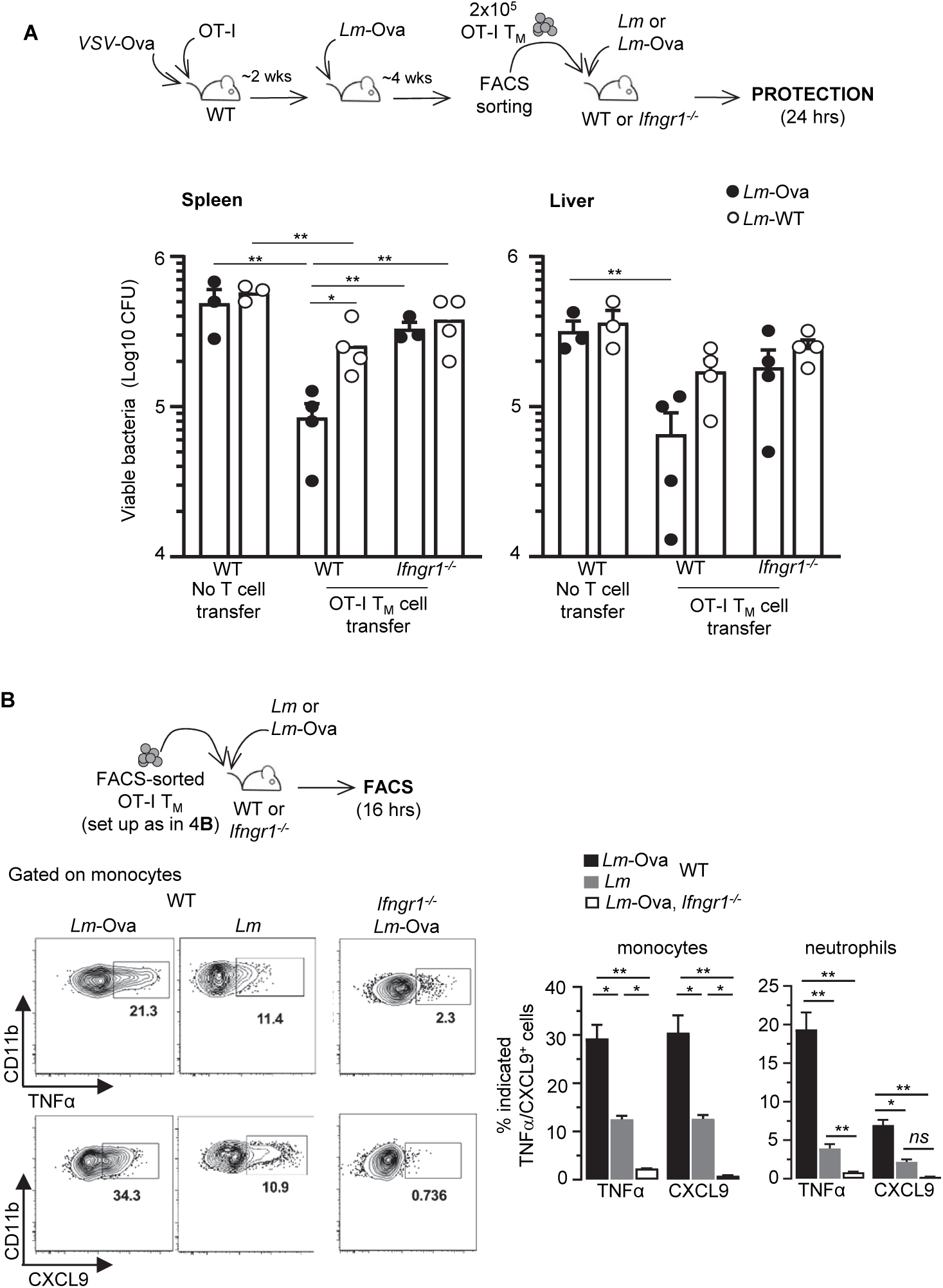
Memory CD8^+^ T cell-mediated protection of vaccinated hosts requires both cognate Ag and IFN_γ_-signaling. 2×10^5^ OT-I T_M_ cells induced using the depicted experimental set up (as in Figure 4B) were transferred in age- and sex-matched WT B6 or *Ifngr1^-/-^* mice, and mice were next challenged with 10^6^ *Lm* or *Lm-*Ova. (A) Control groups did not receive OT-I T_M_ cells. Spleens and livers from challenged mice were harvested 24 hrs later and *Lm* CFUs determining after plating. Bar graphs show 1of 2 representative experiments with each symbol corresponding to 1 individual mouse. (B) Spleens from WT or *Ifngr1^-/-^* mice transferred with OT-I T_M_ cells and challenged with indicated *Lm*, were harvested and cells stained for expression of cell surface CD11b, Ly6C and Ly6G and intracellular TNFα and CXCL9. Representative FACS dot plots are shown and bar graphs pool two representative experiments (n=7 mice) and p- values are indicated.

## Discussion

This study provides an in depth cellular and molecular analysis of how cognate Ag orchestrates the activation of memory CD8^+^ T cells for rapid protection against a recall infection in vaccinated hosts *in vivo*. We reveal that cognate Ag recognition by CD8^+^ T_M_ cells, which occurs selectively on CD11c^hi^ DCs, leads to a much broader gene expression program than inflammation/cytokine-only stimulated counterparts, with multiple pathways affected within only few hours post-stimulation. We also found that IRF4, downstream and proportional to TCR signaling strength, transcriptionally controls the most significantly upregulated cluster of genes in CD8^+^ T_M_ cells that encode for the chemotactic molecules CCL3, CCL4 and XCL1. CD8^+^ T_M_ cell-derived chemokines together with IFNγ then act synergistically to potentiate Ly6C^+^ monocyte antimicrobial effector functions and host protection. This result underlines that chemokines can act as key effector molecules priming innate immune cells, a role distinct from their usual cellular recruitment. Lastly, we demonstrate that this process is spatially restricted to non-diffusive splenic red pulp clusters of Ly6C^+^ monocytes, in which CD8^+^ T_M_ cells arrest upon cognate Ag recognition, to locally deliver activating chemokines and IFNγ to the clustered monocytes, efficiently restraining microbial pathogen spreading and growth.

The current results highlight the importance of rapid microbial pathogen containment, a notion that has been elegantly illustrated in prior reports (Kastenmuller et al., 2013; Sung et al., 2012). First line cellular responders such as splenic marginal zone and LN subcapsular macrophages were reported to rapidly uptake and/or sense microbial pathogens (bacteria, viruses), to subsequently provide chemotactic cues that attract prepositioned memory -but not naïve- CD8^+^ T cells rapidly to the sites of infection. Consistent with these results, intravenously inoculated *Lm* bacteria are rapidly cleared from the blood by marginal zone CD169^+^ macrophages and DCs localized in the splenic RP (Bajenoff et al., 2010; Edelson et al., 2011; Muraille et al., 2005; Neuenhahn et al., 2006; Perez et al., 2017). Following rapid pathogen capture by tissue-resident sentinel cells, a body of evidence suggests that CD8^+^ T_M_ cells home to infectious foci via CXCR3 and/or CCR5 and associated CXCL10, CXCL9 and CCL5 chemokines produced in response to local inflammatory cues such as interferons (Kohlmeier et al., 2010; Kohlmeier et al., 2011; Slutter et al., 2013). A large majority of CD8^+^ T_M_ cells express CXCR3 and CCR5, thus can be readily mobilized for rapid migration, independent from cognate Ag encounter (Maurice et al., 2019). These cells can produce IFNγ in response to cytokines (Kupz et al., 2012; Raue et al., 2013; Soudja et al., 2012), further increasing local chemokine levels in a feedforward positive loop of rapid amplification of the CD8^+^ T_M_ cell response. We report in the current study, that cognate Ag recognition promotes a broad activation program in CD8^+^ T_M_ cells, which includes the early expression of a potent set of chemokines. This finding led us to propose that cognate Ag-triggered CD8^+^ T_M_ cells would amplify the initial chemotactic cues, and act as powerful recruiting orchestrators of both adaptive and innate immune cells, setting the stage for more effective microbial pathogen clearance. Yet and unexpectedly, our results did not support such a model. Rather, we revealed that Ly6C^+^/CCR2^+^ monocytes form clusters in the splenic RP independently from cognate Ag and CD8^+^ T_M_ cells, most likely in response to other infection-driven cues. Adhesion molecules such as ICAM-1, CD11b and CD44, and perhaps not chemotaxis, could be mediating Ly6C^+^ monocyte trafficking to sites of infection as it was shown in the liver of primary *Lm*-infected mice (Shi et al., 2010). Consistent with this idea, we also noted a strong upregulation of ICAM-1 on clustered monocytes in spleen RP. Using IVM imaging, we further revealed that, as expected (Bousso and Robey, 2003; Mempel et al., 2004; Stoll et al., 2002), CD8^+^ T_M_ cells arrest upon cognate Ag recognition, which occurs in monocyte clusters, where they promote their activation through the localized delivery of chemokines and IFNγ. This collectively supports a model where CD8^+^ T_M_ cells only intervene as “late” activators of an already well-coordinated innate immune response, rather than as initial orchestrators of the early steps of that response.

In a previous study using IVM imaging to explore *Lm* infection foci that form in subcapsular DCs (scDCs) of the splenic RP following primary infection (day 5 post-infection), *Lm*-specific effector CD8^+^ T cells were shown to migrate to sites of infection where a mixture of myelomonocytic cells (MMC), that include Ly6C^+^ monocytes and neutrophils, accumulate (Waite et al., 2011). These MMCs drastically reduced blood flow access to the sites of infection and restricted *Lm* growth. *Lm*-specific CD8^+^ T cells were also shown by IVM imaging to undergo both Ag-dependent arrest and Ag-independent reduced motility in the scDC/MMC *Lm*-containing clusters, similarly to our observations. Yet, this study did not address the role that arrested effector CD8^+^ T cells may play in these clusters. Disappearance of *Lm* was associated with effector CD8^+^ T cells regaining motility, but evidence for direct *Lm*-infected killing could not be documented. Together with the large dependence on MMC for *Lm* clearance, these data suggested that, like in the setting of the recall response, non-cytolytic T cell-dependent effector mechanisms were essential. In addition to promoting the local expression of microbicial activities in clustered monocytes, it seems therefore conceivable that the delivery of effector molecules by Ag-arrested CD8^+^ T_M_ cells may also restrict permeability and blood flow in these clusters, to ultimately enhance rapid and effective *Lm* containment and killing.

*Lm* killing and vaccinated host protection during recall infection require TNFα, which Ly6C^+^/CCR2^+^ monocytes are a major source (Nakane et al., 1989; Narni-Mancinelli et al., 2007; Neighbors et al., 2001). TNFα directly triggers microbicidal reactive oxygen species (ROS) both from monocytes and neutrophils and ROS promotes antimicrobial autophagy (Narni-Mancinelli et al., 2011). While we previously showed that IFNγ signaling to Ly6C^+^ monocytes directly induces TNFα production by these cells (Soudja et al., 2014), we now report that both IFNγ signaling and cognate Ag stimulation, and thus chemokine delivery, need to act synergistically to achieve protection of immunized hosts. We also found that IFNγ signaling contributes to chemokine receptor upregulation on monocytes, and robust upregulation of ICAM-1 on the monocytes that altogether may increase cell-cell communication and/or adhesion with T_M_ cells leading to the “sealing” of monocyte clusters for rapid and effective *Lm* clearance. While we did not monitor neutrophil dynamics here, neutrophils are well known to undergo massive recruitment and activation in infected spleens, and we and others have shown previously that they cluster with Ly6C^+^ monocytes and CD8^+^ T_M_ cells at infection foci (Alexandre et al., 2015; Bajenoff et al., 2010; Soudja et al., 2014). However, and in contrast to Ly6C^+^ monocytes, neutrophils neither express nor upregulate high levels of CCR1 and CCR5, suggesting that CD8^+^ T_M_ cell-derived chemokines are unlikely to account for activating neutrophils in this setting. Fine-tuning of monocyte activation in response to local chemokine levels may however regulate their secretion of TNFα, which directly promotes ROS production and pathogen killing.

Another important finding in our study relates to the rapid, transcriptionally controlled and coordinated production of CCL3, CCL4 and XCL1 chemokines by CD8^+^ T_M_ cells induced upon vaccination with both *Lm* and *VSV* in response to cognate Ag recognition. These results are consistent with two recent reports that utilized multiple models of acute and chronic infections, as well as *ex vivo* stimulation assays, and outline that the robust chemokine signature is a key and important feature of both Ag-stimulated effector and memory CD8^+^ T cells (Davenport et al., 2020; Eberlein et al., 2020). CD8^+^ T_M_ cells undergoing repetitive *in vivo* stimulations were also reported to significantly upregulate genes encoding for these chemokines (Wirth et al., 2010). Our study further reveal that CCL3, CCL4 and XCL1 chemokines produced by cognate Ag-stimulated CD8^+^ T_M_ cells, are directly under the control of the IRF4 transcriptional regulator, a known amplifying rheostat downstream of TCR signaling (Man et al., 2013). IRF4 may indeed enable the graded production of these chemokines by CD8^+^ T_M_ cells, proportionally to the strength of TCR signaling, and this could represent a mechanism to limit tissue-associated damages, when weak epitopes are presented to Ag-specific CD8^+^ T_M_ cells. In the context of strong epitope stimulation, however, our results suggest that chemokines are secreted concomitantly to CD8^+^ T_M_ cells arrest in monocyte clusters, promoting their increased production of TNFα both *in vitro* and *in vivo*. The fact that both IFNγ signaling and cognate Ag recognition, are required for vaccinated host protection is consistent with a key role for chemokines in potentiating clustered monocyte antimicrobial functions. These findings also highlight that chemokines can prime innate immune cell effector functions, clearly delineating a role distinct from usual chemotaxis.

While our study focuses on systemic and SLO-derived memory CD8^+^ T cell responses, multiple evidence suggest that the current mechanisms are also relevant in the context of tissue-resident memory CD8^+^ T cell responses. In several models of viral infection (skin, vagina, lung), T_RM_ cells -both CD8^+^ and CD4^+^- quickly initiate and orchestrate a rapid mucosal response upon cognate Ag encounter, through local production of IFNγ and subsequent CXCL9 (Ariotti et al., 2014; Beura et al., 2018; Iijima and Iwasaki, 2014; Kohlmeier et al., 2009; Kohlmeier et al., 2010; Schenkel et al., 2014). As discussed earlier, CXCL9 enables migration of more circulating T_M_ cells to sites of infection, enhancing the activation of local DCs and NK cells and the establishment of an IFNγ-driven antiviral state providing broad protective immunity against unrelated microbial pathogens. In these studies, reactivation of CD8^+^ T_M_ cells and the production of activating IFNγ required cognate Ag recognition, yet many reports monitoring systemic CD8^+^ T_M_ cells, have also established that CD8^+^ T_M_ cells in SLO undergo cytokine-mediated activation (Bedoui et al., 2009; Berg et al., 2003; Maurice et al., 2019; Raue et al., 2013; Soudja et al., 2012). This seemingly discrepant result may be a reflection of tissue-specific mechanisms. In fact, it was recently shown that LN CD8^+^ T_M_ cells strictly require cognate Ag to be presented by XCR1^+^ DCs while lung T_RM_ can be reactivated by both hematopoietic and non-hematopoietic cells (Low et al., 2020). Here, cognate Ag presentation by hematopoietic versus non-hematopoietic-derived cells to CD8^+^ T_RM_ cells was also proposed to dictate distinct functional outcomes with hematopoietic-derived APCs restraining an excessive inflammatory program in CD8^+^ T_RM_ cells, presumably as a safeguard mechanism against collateral tissue damages. Interestingly, non-hematopoietic Ag presentation was associated with a proliferative program, and largely prevented cytokine-mediated activation of CD8^+^ T_RM_ cells. It is noteworthy this study used the Nur77^GFP^ reporter system a read out of TCR-dependent cognate Ag stimulation, thereby only focusing on early Ag-dependent CD8^+^ T_M_ cell-expression programs. Other reports using complex biological read-outs (e.g., proliferation, protection) have supported a more prominent role for recruited or tissue-resident DCs in the reactivation of CD8^+^ T_RM_ cells, raising the possibility that different memory cell-intrinsic mechanisms of regulation may be specifically programmed upon DC- versus non-hematopoietic cell-mediated activation (Shin et al., 2016; Wakim et al., 2008).

In conclusion, this work provides a detailed analysis of how cognate Ag stimulation induces early changes in systemic CD8^+^ T cell memory, and how these changes enable the rapid control of microbial pathogen in vaccinated hosts *in situ*. Perhaps contrasting with the widely accepted view, our results suggest that memory CD8^+^ T cells are not essential in orchestrating the initial immune response. The initiating response is largely regulated by tissue-specific cues and innate immune cells before CD8^+^ T_M_ cells intervene to make the innate cellular effector response highly effective. The findings also reveal that a combination of cells (T_M_ cells, monocytes, neutrophils, DCs) and signals (Ag, IFNγ, chemokines) are needed to achieve early protection, which suggests many levels of fine tuning are possible to make such response the most efficient and less damaging to vaccinated host.

## EXPERIMENTAL PROCEDURES

### Ethics Statement

This study was carried out in strict accordance with the recommendations by the animal use committee at the Albert Einstein College of Medicine. All efforts were made to minimize suffering and provide humane treatment to the animals included in the study.

### Mice

All mice were bred in our SPF animal facility at the Albert Einstein College of Medicine. We used wild-type (WT) C57BL/6J (B6) 6-8 weeks old male or female mice, congenic CD45.1^+/+^ (JAX#002014), B6-K^d^ (Lauvau et al., 2001), OT-I^+^ (JAX#003831) crossed to *Rosa26-Actin-tomato-stop^loxP/loxP(or fl/fl)^*^-GFP^ (Td^+^)(JAX#007576), gBT-I^+^ (kind gift Francis Carbone (Mueller et al., 2002)) crossed to *UBC^GFP/GFP^* (JAX#004353) or to CD45.1^+/+^ mice, *Ccr2*^DTR-CFP/WT^ (kind gift Eric Pamer (Hohl et al., 2009)), *Itgax/Cd11c*^DTR/WT^ (JAX#004509), *Rosa26^Cre-ERT2^* (JAX#008463), *Irf4^loxP/loxP (or fl/fl)^* (JAX#009380), *Ifngr1^-/-^* (JAX#003288) purchased from the Jackson labs unless otherwise indicated. All mice are on the B6 genetic background unless otherwise specified.

### Microbial pathogens and mouse infections

*Listeria monocytogenes (Lm):* Mice were inoculated with *Lm*, *Lm* expressing the ovalbumin (*Lm*-Ova, kind gift H. Shen, U Penn) or Ova and *Herpes Simplex Virus 2 (HSV-2)* glycoprotein B 498-505 epitope (*Lm*-Ova-gB, kind gift D. Zehn, TUM), all expressed under the LLO/*Hly* promoter. All *Lm* were prepared after passaging into WT B6 mice, by growing to log phase (OD_600_∼0.3-0.4) and kept as frozen aliquots for single use in -80°C. For infections, bacteria were grown to a logarithmic phase (OD600∼0.05-0.15) in broth heart infusion medium, diluted in PBS to infecting concentration and injected intravenously (i.v). We used 0.1×LD_50_, i.e., 10^4^ *Lm* CFUs for primary immunizations, and 10^6^ *Lm* CFUs for secondary challenge infections (∼6 wks later). All *Lm* are on the 10403s genetic background.

*Vesicular Stomatitis Virus* (*VSV*): Single use frozen aliquots of *VSV* encoding Ova (*VSV*-Ova, gift Kamal Khanna, NYU) kept at -80°C were thawed and diluted in cold PBS right before mouse primary i.v. infections with 2×10^5^ PFUs. For secondary challenge infections of *VSV*- immunized mice (∼6 wks later), we used 10^6^ *Lm* CFUs.

### Preparation of cell suspensions for flow cytometry and adoptive transfers

Spleens were dissociated on a nylon mesh and lysed in red blood cells (RBC) lysis buffer (0.83% NH_4_Cl vol/vol), prior to incubation in HBSS medium with 4,000 U/mL collagenase I and 0.1 mg/mL DNase I. BM cells were obtained by flushing femur with complete medium (RPMI 1640, 10% FBS, 1% Penicillin/Streptomycin, 55μM β-mercaptoethanol, 1mM Sodium Pyruvate, 1X Glutamax, 1X non-essential amino acids) containing 10% FCS.

### Cell-staining for FACS analysis

Cell suspensions were incubated with 2.4G2 Fc Block and stained in PBS 1%FCS, 2mM EDTA, 0.02% sodium azide with fluorescently tagged Abs purchased from eBioscience, BD Biosciences, R&D systems, or BioLegend (See Table S3) or in same instance tetramers. For tetramers, biotinylated monomers (1 mg/mL) obtained from the NIH tetramer Core Facility were conjugated with PE-labeled Streptavidin (1mg/mL) as follow: 6.4 μL of PE-Streptavidin were added to 10 μL of monomers every 15 min 4 times on ice. Newly generated tetramers (1/400- 1/500 dilution) were then used to stain spleen cells for 1 hr at 4℃. For IRF4 transcription factor (TF) intracellular staining (IRF4), cells were fixed in eBioscience Fixation/Permeabilization buffer prior to anti-IRF4 mAb staining in eBioscience Permeabilization buffer for 30 min. For intracellular cytokine staining (ICS), cells were first incubated for 4 hrs at 37°C/5%CO_2_ in complete medium 10% FCS, with Golgi Plug/Golgi Stop. Next, cells were stained for cell-surface marker expression and fixed in IC fixation buffer (eBioscience) prior to permeabilization ∼1h in presence of Abs against intracellular cytokines (IFNγ, TNFα) and chemokines (CXCL9, CCL3, CCL4, XCL1). For CCL3, CCL4 and XCL1, a Donkey Anti-Goat (DAG, 2µg/mL) secondary Ab was used. Data acquisition was done using a FACSAria III flow cytometer. All flow cytometry data were analyzed using FlowJo v9 software (TreeStar).

### Adoptive T cell and monocyte transfers

#### T cell transfers

For naïve OT-I and gBT-I cells, WT mice were adoptively transferred with ∼1,000 OT-I Td^+^ and 50,000 gBT-I cells isolated from the spleen of OT-I Td^+^ and gBT-I CD45.1^+/+^ mice. The next day, mice were immunized with *Lm*-Ova-gB. Immunized mice were next used ∼6 wks later to investigate OT-I and gBT-I T_M_ cell reactivation by FACS and IVM. For adoptive transfers of OT-I memory cells, WT mice were first adoptively transferred with ∼1,000 naive OT-I Td^+^ cells as above, immunized the next day with *VSV*-Ova and challenged 2 wks later with 10^4^ *Lm*-Ova. After ∼4 wks, spleens were harvested and CD8^+^ T cells CD8^+^ T cells were negatively selected from spleen using anti-CD4, anti-CD11b, anti-MHC II, anti-TER119, anti-B220 and anti-CD19 mAbs (Table S3), which all were added and incubated at 5 μg/mL for 30 min at 4C. Cells were then washed and incubated with anti-rat Ab magnetic beads at 1 bead/target cell for 40 min at 4C (Dynabeads sheep anti-rat IgG, Invitrogen). Cells were sorted into 3mL of complete media (RPMI 1640, 10% FBS, 1% Penicillin/Streptomycin, 55μM β-mercaptoethanol, 1mM Sodium Pyruvate, 1X Glutamax, 1X non-essential amino acids) using a 4 laser BD FACS Aria III cell sorter. 2×10^5^ OT-I T_M_ cells were transferred to either WT, *Ccr2^CFP^* or *Ifngr1^-/-^* recipient mice further challenged with 10^6^ *Lm* or *Lm*-Ova for analysis of memory functions, protection or IVM.

#### Monocyte transfers for chemokine receptor expression analysis

5×10^6^ BM cells from WT CD45.1^+/-^ and *Ifngr1^-/-^* CD45.2^+/+^ donors were co- transferred to mice immunized with *Lm*-Ova 6 wks before, and further challenged or not with *Lm*. Spleen cells were next stained for chemokine receptor expression on monocytes from BM donor derived cells.

### Generation of bone-marrow chimera mice

Lethally irradiated (1,200 rads) B6-K^d^ mice were immediately reconstituted with a total of 2×10^6^ BM cells isolated from i) C*cr2^DTR^*K^d^ and K^d^, ii) C*cr2^DTR^*K^d^ and WT, iii) C*d11c^DTR^*K^d^ and WT, iv) K^d^ and WT mice, at a 7:3 ratio, respectively. Donor BM cells were depleted of CD8 and CD4 T cells from WT BM cells using anti-CD8β (clone H35) and anti-CD4 (clone GK1.5) mAbs prior to reconstitution. Chimerism of reconstituted mice was checked ∼6 wks later in the blood, prior to immunizations.

### *In vivo* treatments

#### Monocytes and DCs depletion

CCR2^+^ and CD11c^+^ cells were depleted from diphtheria toxin receptor (DTR)-expressing mice upon intraperitoneal (i.p.) injection of 10 ng/g of mouse body weight of diphtheria toxin (DT, Calbiochem) 12 hrs prior to *Lm* challenge infection.

#### *Irf4* depletion

4-hydroxytamoxifen (Tx) (4-OHT, #T5648, Sigma-Aldrich) was dissolved in sunflower oil to a concentration of 10 mg/mL for i.p. injection. 3,000 OT-I-*Rosa26- Cre^ERT2^Irf4^fl/fl^* and 1,000 OT-I *Irf4*^WT^ were co-transferred to WT B6 mice which were immunized with *Lm*-Ova the next day. Six weeks later, and prior to challenge, mice were treated with Tx (100 µL, 1mg/injection) or vehicle (100µL of sunflower oil) for 5 days and 24 hrs after the last Tx injection, mice were challenged with *Lm*-Ova.

#### Ly6C-PE antibody for monocyte staining *in situ*

For IVM analysis, 10µg of Ly6C-PE (Clone HK1.4, Rat IgG2a, Biolegend) mAb was inoculated to mice i.v. at the time of *Lm* challenge or 1 hr before sacrificing mice. For FACS analysis, 5µg of Ly6C-PE mAb was injected to challenged mice 1hr prior to the sacrifice.

### *In vitro* activation assays

#### Quantification of CCL3 and IFN-γ secretion

10^6^ splenocytes from mice immunized with *Lm*- Ova and challenged or not 6 wks later with *Lm*-Ova for 16 hrs, were incubated in 96-flat bottom wells with complete medium only or in presence of Golgi Plug/Stop for 4 hrs at 37°C. CCL3 (Thermofisher) and IFN-γ (Biolegend) production in culture supernatants was then quantified by ELISA.

#### Measure of chemokines expression by *ex vivo* restimulated OT-I T_M_ cells

Splenocytes from mice immunized with *Lm-*Ova 6 wks prior were co-incubated with SIINFEKL peptide (10^8^M) with Golgi Plug/Stop and i) with or without cycloheximide (Translation inhibitor, 10µg/mL, Sigma-Aldrich) or Actinomycin D (Transcription inhibitor, 8µM, Sigma-Aldrich) for 1, 2, 3 and 4 hrs in complete medium at 37°C ; ii) with the SCG-CBP30 IRF4 inhibitor (Selleckchem, 20µM). Cells were next stained as described above including for intracellular expression of CCL3 and/or CCL4 and XCL1.

#### Measure of TNFα expression in *ex vivo* stimulated monocytes and neutrophils

10^4^ monocytes or neutrophils FACS-sorted (Aria III) from mice primary immunized and challenged with *Lm*-Ova 6 wks later for 16 hrs were co-incubated in 96-round bottom wells and complete medium, with HK*Lm*, rCCL3, rCCL4, rXCL1 or the combination of the three recombinant chemokines, in presence of Golgi Plug/Stop for 4 hrs at 37°C before staining for cell surface and intracellular markers for FACS analysis. In CCR5 and CCR1 blocking experiments, cells were incubated with both CCR5 (1µM, Maraviroc, Cayman Chemical Company) and CCR1 (1µM, J113863, Santa Cruz) chemical inhibitors or control DMSO 30 min prior to adding recombinant chemokines or HK*Lm*.

### RNA-Sequencing

#### Samples and library preparation

1,000 OT-I Td^+^ and 1,000 gBT-I CD45.1^+/+^ T_M_ cells were flow-purified (FACS Aria III) following the same procedure as for the adoptive T_M_ cell transfers described before, from 6 wks *Lm*-Ova-gB-immunized mice either left unchallenged or challenged with *Lm*-Ova. T_M_ cells were directly sorted into 1X lysis buffer and cDNA was synthesized and amplified directly from intact cells using SMART-Seq v4 Ultra Low Input RNA Kit for Sequencing (Takara Bio USA) according to the manufacturer protocol. cDNA was isolated using the Agencourt AMPure XP Kit (Beckman Coulter, Brea, CA) and quantified using the Qubit dsDNA High Sensitivity Assay Kit (Life Invitrogen) on an Agilent 2100 Bioanalyser (Agilent Technologies, Santa Clara, CA). The library preparation was performed using the Nextera XT DNA Library Preparation Kit (Illumina Inc., San Diego, CA). Samples were sequenced to depths of up to 16.7 million single-end 75 nt length reads per sample using the Illumina NextSeq 500/550 High Output v2 kit (75 cycles) on an Illumina NextSeq 500 Sequencing System. Image analysis, base calling, and generation of sequence reads were produced using the NextSeq Control Software v2.0 (NCS) and Real-Time Analysis Software v2 (RTA). Data was converted to FASTQ files using the bcl2fastq2 v2.20 software (Illumina Inc.). Sequencing data was initially quality-checked using FastQC, before alignment and initial analysis. Reads were aligned to the Mouse reference mm10 using STAR aligner (v2.4.2a) (Dobin et al., 2013). Quantification of genes annotated in Gencode vM5 were performed using featureCounts (v1.4.3) and quantification of transcripts using Kalisto (Bray et al., 2016). QC was collected with Picard (v1.83) and RSeQC (Wang et al., 2012)(http://broadinstitute.github.io/picard/). Normalization of feature counts was done using the DESeq2 package, version 1.10.1. Prior to analysis, non-relevant batch effect (such as library preparation or sequencing batch) was identified using unsupervised principal component analysis (PCA) and analysis was corrected for batch effects through our model. Differentially expressed genes were identify using negative binomial distribution as implemented in DESeq 2 (R package (Love et al., 2014)). Significantly up and down regulated genes (DEG) were defined with an FDR step-up *p* ≤ 0.05 and a fold-change ≥±1.5. The raw data from the NCBI database (GEO GSE160280) was subsequently analysed for enrichment of GO terms and the KEGG pathways, implemented in the clusterprofiler (R package, function enrichGO or enrichKEGG) ; a pathway is considered significantly-enriched if the enrichment score is ≥1.5 (equivalent to a *p* ≤ 0.05).

### Intravital and Explant Imaging

For intravital imaging, mice were anesthetized with isoflurane and the spleen was surgically exposed and elevated above the body of the mouse. A glass coverslip was carefully applied to the top of the spleen to create an imaging window. Mice were kept at 37°C using a custom heating platform. Imaging was performed on an Olympus FVE-1200 upright microscope using a 25X 1.04 NA objective and a Deepsee MaiTai Ti-Sapphire pulsed laser (Spectra Physics) tuned to 870nm. To maintain temperature and limit infiltrating light, the microscope was fitted with a custom-built incubator chamber heated to 37°C. 512×512 Z-stack images were acquired every 60 seconds with 5 um steps. For explant imaging, mice were euthanized with CO_2_ and spleens immediately harvested. Spleens were affixed to coverglass using VetBond (3M) on the medulla. Tiled images were acquired using 320×320 Z-stack images with 15 um steps. Tiled images were stitched using Olympus Fluoview Software. Cell tracking, drift correction, and monocyte volume analysis was carried out using Imaris 9.2 (Bitplane).

### Measure of protective immunity

2×10^5^ OT-I T_M_ cells were flow-sorted as described before and adoptively transferred to WT or *Ifngr1^-/-^* mice and the next day challenged with *Lm* or *Lm*-Ova. Untransferred mice were used as control groups. To measure *Lm* titers, spleen and liver were harvested 24 hrs post challenge infection and dissociated on metal screens in 10 mL of water/0.1% Triton X-100 (Sigma-Aldrich). Serial dilutions were performed in the same buffer, and 100 μl were plated onto BHI media plates. *Lm* CFU numbers were counted 24 hrs later.

### Statistics

Statistical significance was calculated using an unpaired Student t test with GraphPad Prism software and two-tailed p values are given as: (*) p<0.1; (**) p<0.01; (***) p<0.001; (****) p<0.0001 and (ns) p>0.1. All p values of 0.05 or less were considered significant and are referred to as such in the text.

## Acknowledgments

We thank the Einstein FACS and genomic core facilities.

## Author contribution

MB designed, performed and interpreted all experiments and assembled all Figures, and wrote the manuscript with GL. ZB and DF designed, conducted analyzed and interpreted IVM experiments and contributed to scientific discussions and manuscript editing. FD with MB and GL analyzed and interpreted all transcriptomic data. EG and CK contributed to multiple mouse experiments with MB. SS generated the *Lm*-LLO_Ser92_-Ova strain. GL designed and interpreted experiments with MB and all other authors, contributed to Figure design and editing, and wrote the paper with MB.

## Data materials and availability

The accession number for the microarrays and RNA-seq data reported in this paper is GEO:GSE160280. All data is available in the main text or the supplementary materials.

## Supplemental Figure Legends

**Figure S1, related to Figure 2.**
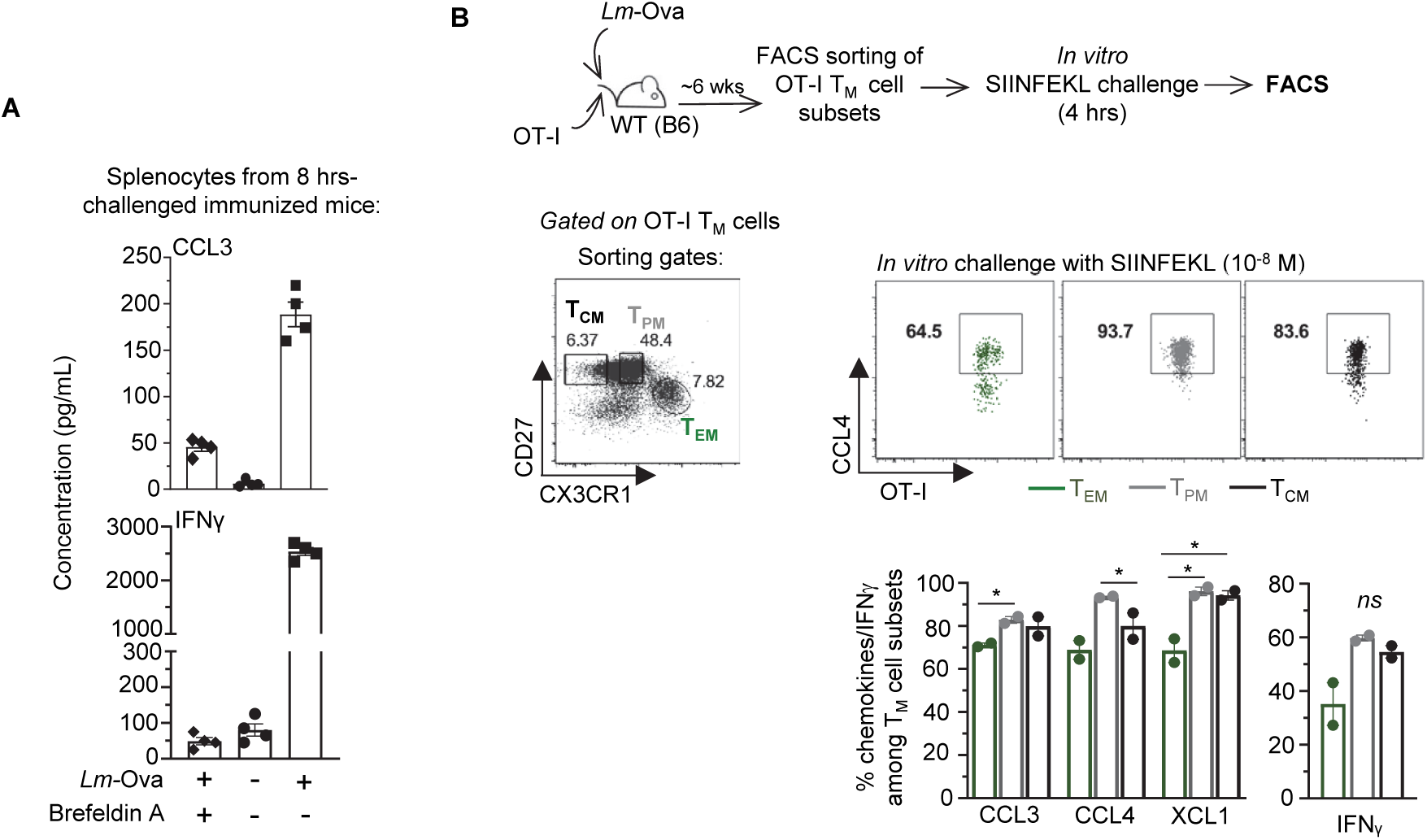
(A) Spleen cells were isolated from *Lm*-Ova-immunized mice (10^4^) that were rechallenged 6 wks later with 10^6^ *Lm*-Ova. At 8 hrs post challenge, spleen cells were incubated or not with Golgi Plug/Stop, before collecting supernatants 4 hrs later. CCL3 and IFNγ were next quantified by ELISA. (B) Subsets of OT-I T_M_ subsets were flow-sorted from the spleens of mice transferred with naïve OT-I cells and immunized with 10^4^ *Lm*-Ova 6 wks before (schematic), based on CX3CR1 and CD27 cell surface marker expression (T_EM_: CX3CR1^hi^CD27^low^, T_PM_: CX3CR1^int^CD27^hi^, T_CM_: CX3CR1^low^CD27^hi^). Sorted (gates are shown) OT-I T_M_ subsets were next stimulated for 4 hrs with the SIINFEKL peptide (10^-8^M) *in vitro* and stained for cell-surface CD8, CXCR3, KLRG1 and intracellular CCL3, CCL4, XCL1 and IFNL. Representative dot plots with summary bar graphs (each symbol is 1 mouse) show the proportion of chemokines^+^ or cytokines^+^ cells among CD8^+^ T_M_ subsets with indicated p-value.

**Figure S2, related to Figure 4.**
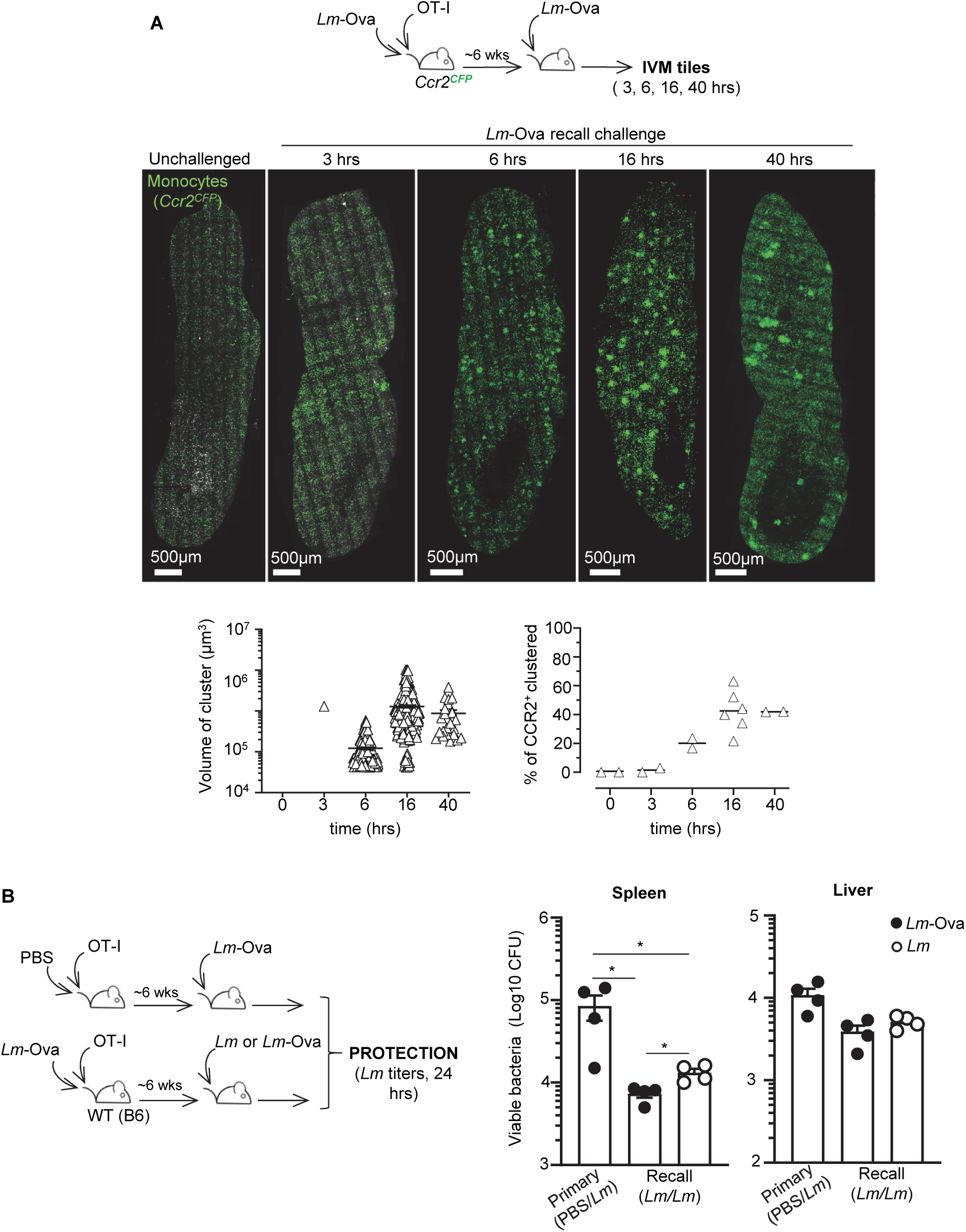
(A) *Ccr2^CFP^* mice grafted with OT-I cells were immunized with 10^4^ *Lm*-Ova and 6 wks later challenged or not with 10^6^ *Lm*-Ova for 3, 6, 16 and 40 hrs. Ccr2^CFP^ monocytes are in green and representative mouse spleen tile reconstructions are shown. Graphs shows the volume of individual clusters and the average proportion of CCR2^+^ monocytes clustered in a pool of 2 independent experiments (n=2-5 mice). (B) Age- and sex-matched WT B6 mice were transferred with OT-I cells and injected with PBS or immunized with 10^4^ *Lm-*Ova. 6 wks later, PBS-injected mice were challenged with 10^6^ *Lm-*Ova (“Primary”), and *Lm-*Ova-immunized mice were challenged with 10^6^ *Lm* or *Lm-*Ova (“Recall”). Bacterial titers in spleens and livers were determined 24hrs post challenge. Bar graphs represent 1 of 2 representative experiments with each symbol corresponding to an individual mouse (n=4 mice).

**Figure S3, related to Figure 4.**
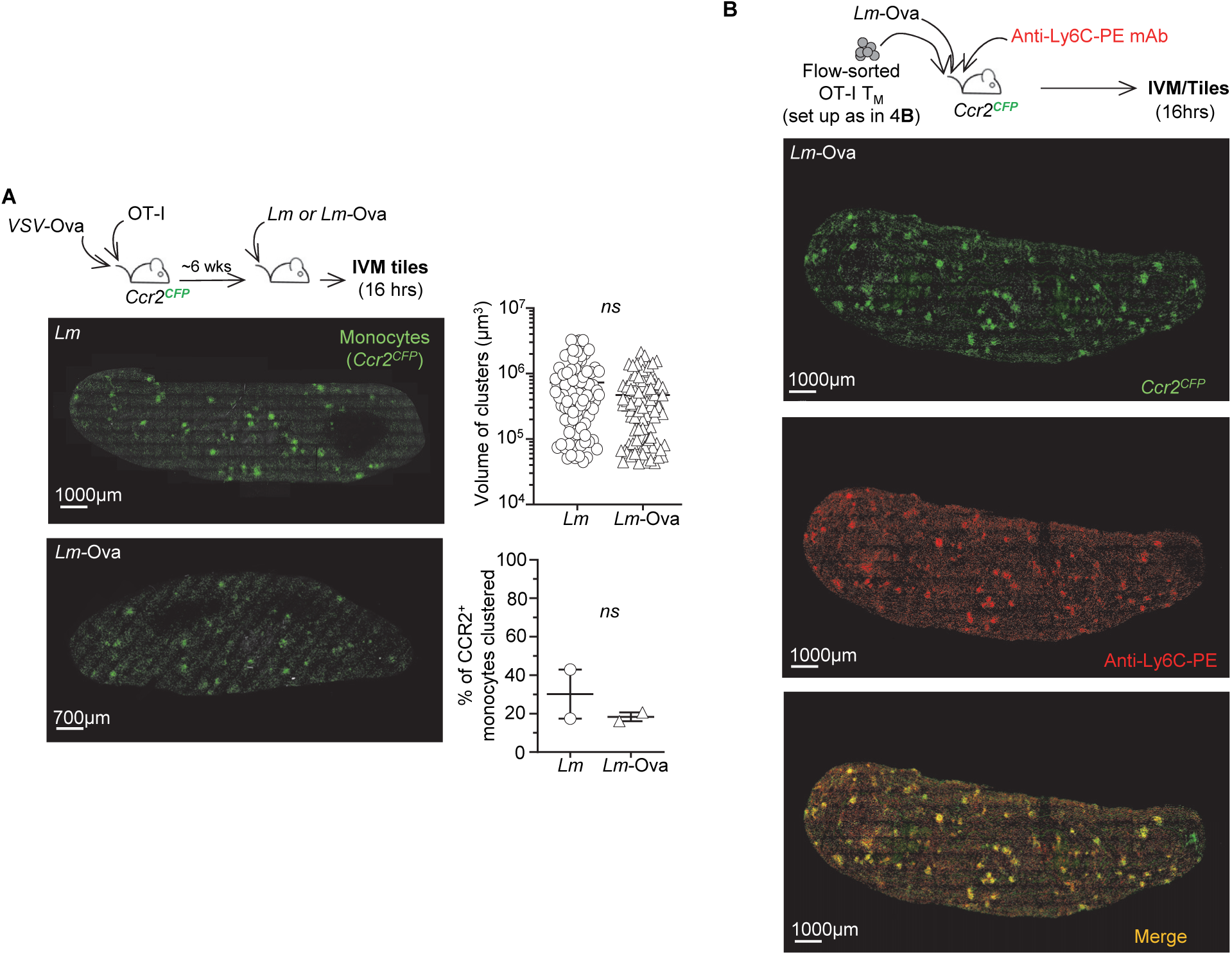
(A) *Ccr2^CFP^* mice grafted with OT-I cells were immunized with *VSV*-Ova (2×10^5^ PFU) and 6 wks later challenged with 10^6^ *Lm* or *Lm*-Ova for 16 hrs. Ccr2^CFP^ monocytes are in green and representative tiles of reconstructed mouse spleens are shown. Graphs show the volume of individual clusters and the average proportion of CCR2^+^ monocytes clustered in 2 independent replicate experiments (n=2 mice). (B) 2×10^5^ flow-sorted- OT-I T_M_ cells generated as depicted in Figure 4B, were transferred to naive *Ccr2^CFP^* that were also injected with Ly6C-PE mAb (10µg) and 10^6^ *Lm*-Ova for 16 hrs. Representative tiles of reconstructed mouse spleens in 1 of 2 replicate experiments are shown with Ccr2^CFP^ monocytes in green and Ly6C-PE^+^ monocytes in red (n=4 mice). Green and red signals are merged in yellow.

**Figure S4, related to Figure 5.**
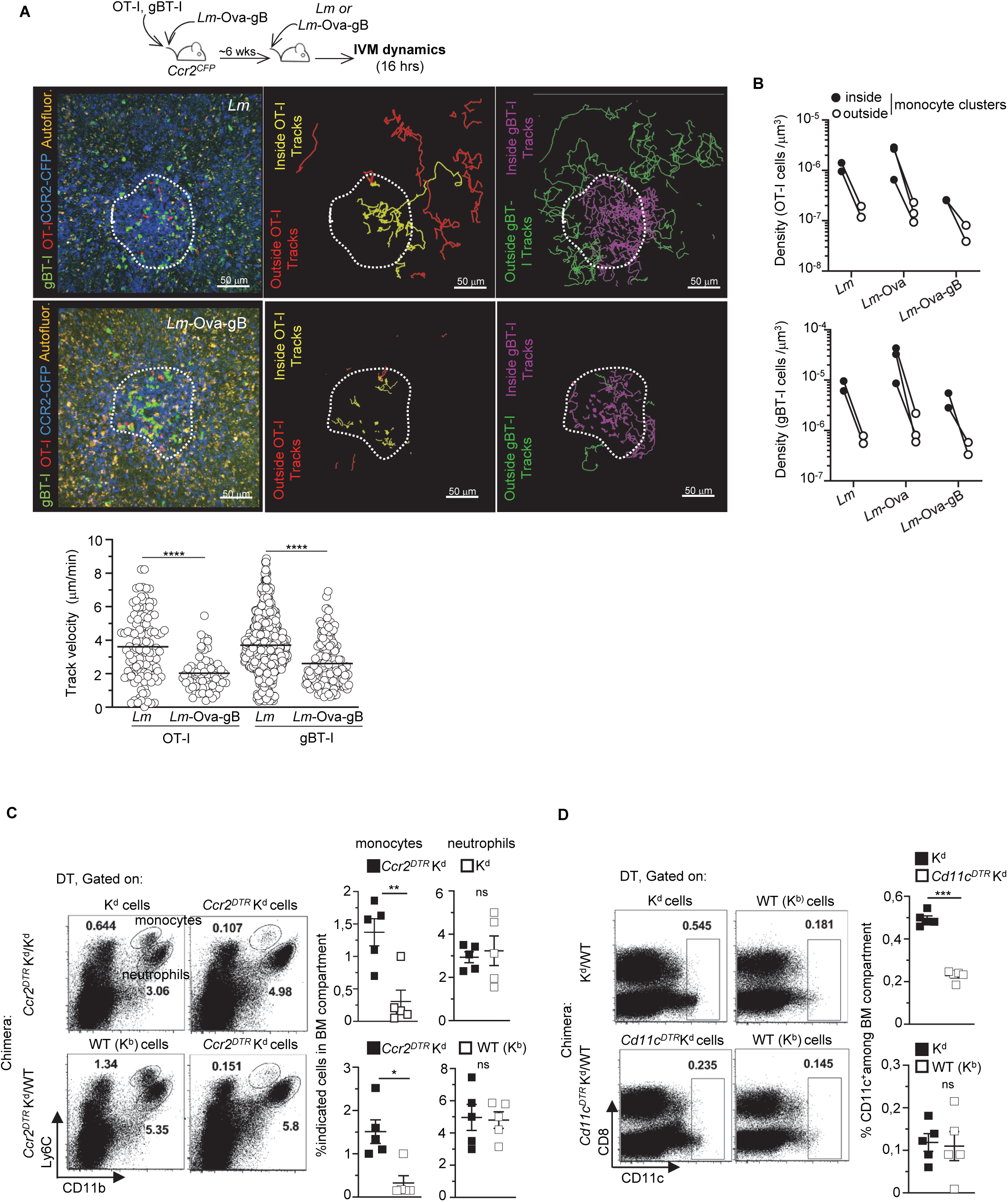
*Ccr2^CFP^* mice were co-transferred with naïve OT-I Td^+^ and gBT-I GFP^+^ cells and immunized with 10^4^ *Lm-*Ova-gB. Six wks later, mice were challenged with 10^6^ *Lm* or *Lm-*Ova-gB and IVM images in the spleen of live mice were recorded 16 hrs later. (A) Representative IVM image (right) of OT-I (red) and gBT-I (green) T_M_ cells localized in a cluster (delimited area by white dashed line) of CCR2^CFP^ monocytes (blue) are shown with autofluorescence (yellow). OT-I T_M_ (outside, red and inside, yellow) and gBT-I T_M_ cell tracks (outside, green and inside, purple) inside and outside the cluster of CCR2^CFP^ monocytes are also shown (center and left images) . Graphs represent the speeds of OT-I and gBT-I T_M_ cells in the clusters. (B) Graph represent the density of OT-I or gBT-I T_M_ cells inside and outside of monocyteclusters after *Lm, Lm-*Ova or *Lm*-Ova-gB challenges. (C, D) Efficiency of DT-mediated depletion in indicated groups and compartments of mixed BM chimeras. Representative FACS dot plots in a pool of 2 experiments with p-value are shown (n=5 mice).

**Figure S5, related to Figure 6.**
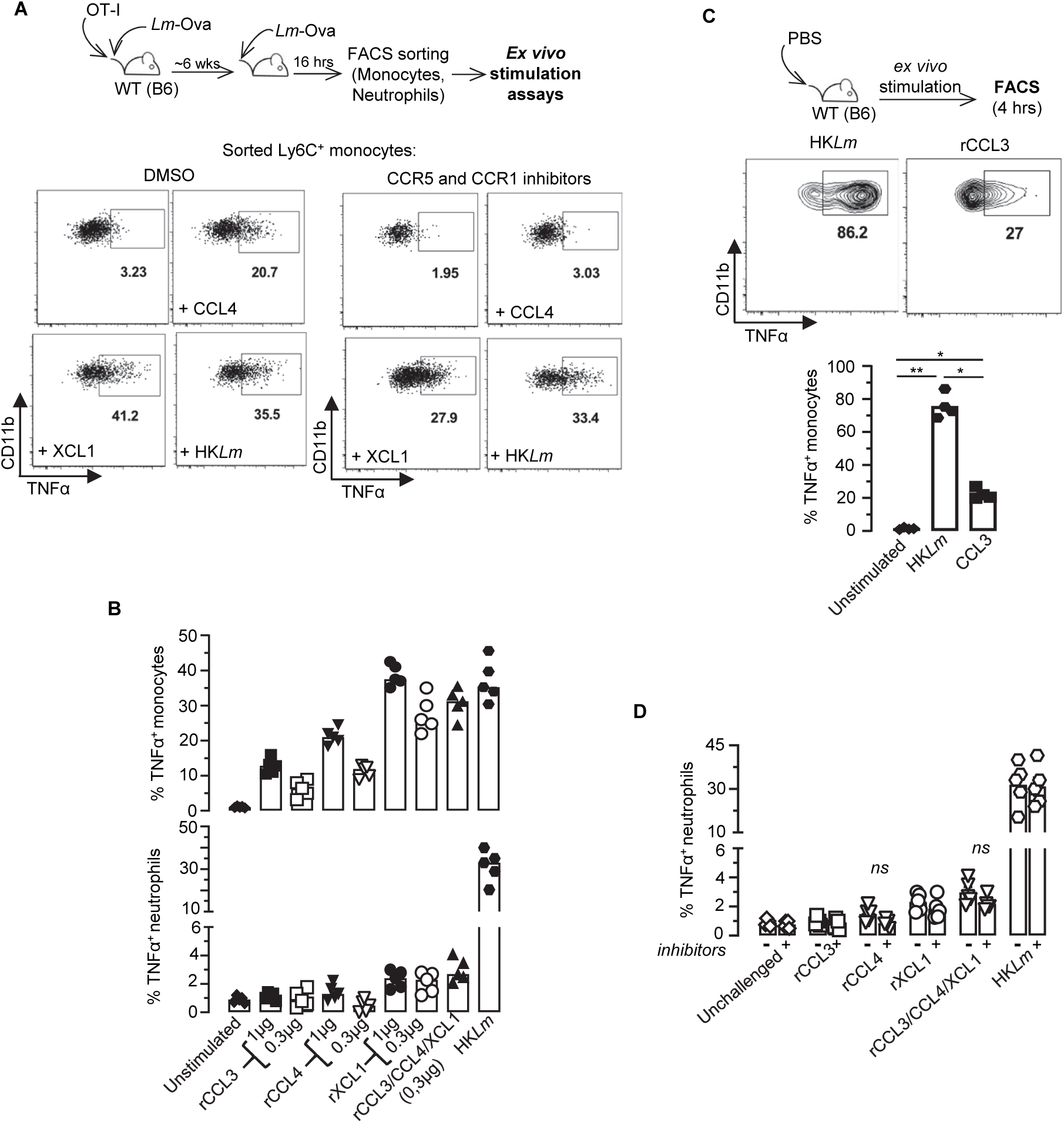
(A, B) Mice grafted with OT-I cells were immunized with *Lm*- Ova and 6 wks later challenged with 10^6^ *Lm-*Ova. Sixteen hrs post-challenge, monocytes and neutrophils were sorted from spleen and stimulated for 4 hrs with or without recombinant chemokines at indicated concentrations, or with HK*Lm* and with or without CCR1/5 inhibitors (depicted in Figure 6C). Cells were next stained for cell surface expression of CD11b, Ly6C, Ly6G and intracellular TNFα. Representative FACS dot plots are shown. (B) Graphs show TNFα ^+^ monocytes and neutrophils frequency after 4hrs incubation with recombinant chemokines (1 and 0.3µg) or HK*Lm*. Bar graphs (each symbol is 1 mouse) represent the pool of 2 independent replicate experiments with p-values indicated. (C) Splenocytes from WT naive mice injected with PBS were stimulated *in vitro* with HK*Lm* or rCCL3 (1µg). Cells were stained as above. Representative FACS dot plots with summary bar graphs (each symbol is 1 mouse) with indicated p-value are shown. (D) TNFα expression in flow-sorted neutrophils as depicted in Figure 6C, incubated with CCR5 and CCR1 inhibitors or DMSO vehicle. Bar graphs (each symbol is 1 mouse) represent the pool of 2 independent replicate experiments with p-values indicated.

**Figure S6, related to Figure 6.**
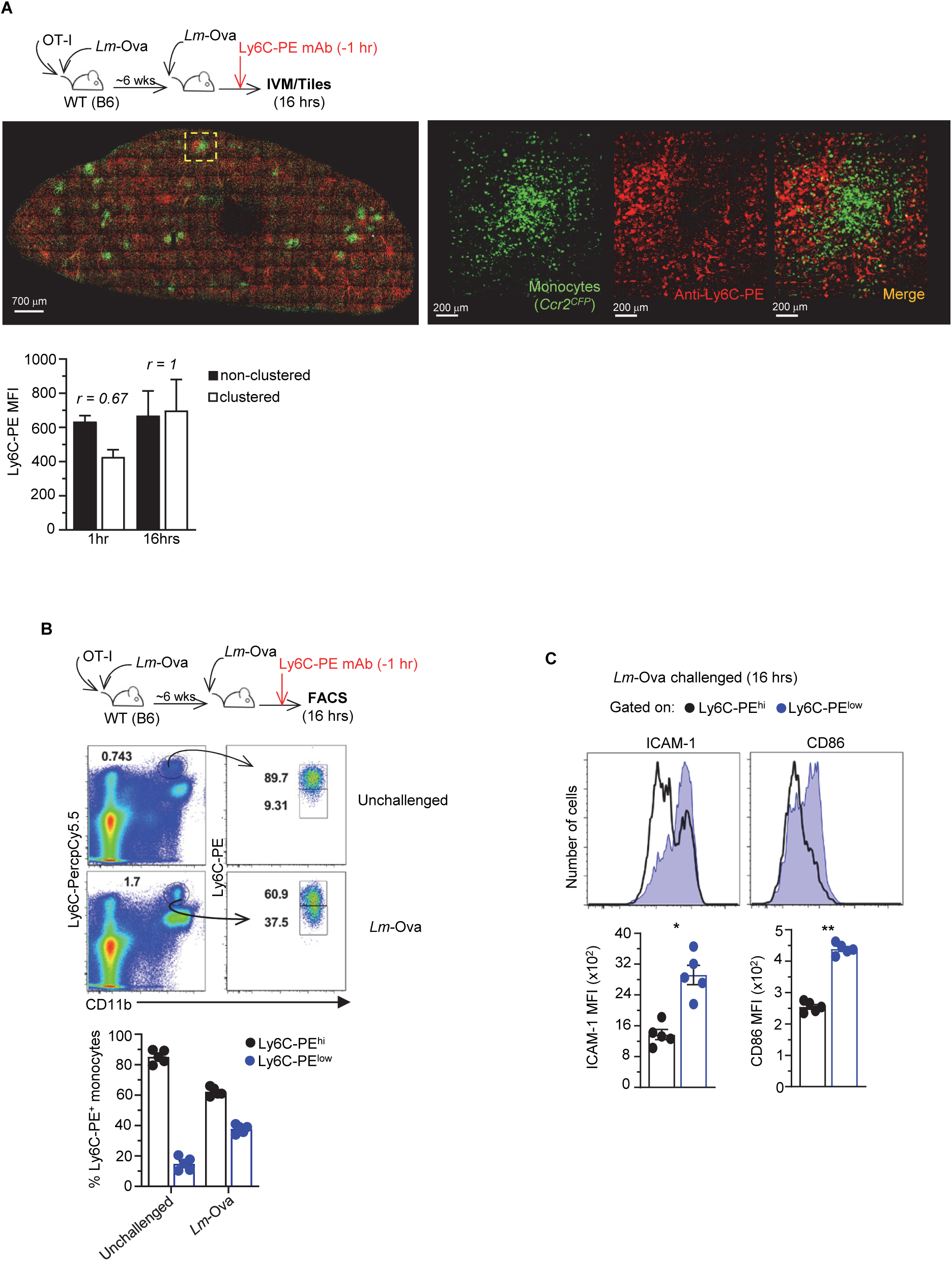
(A-B) *Lm*-Ova-immunized mice were co-transferred with naive OT-I Td^+^ cells and challenged or not ∼6 wks later with *Lm*-Ova for 16 hrs. 1hr before sacrifice, mice were injected i.v. with Ly6C-PE mAb. (A) Representative tiles of reconstructed mouse spleens in 1 of 2 replicate experiments are shown with Ccr2^CFP^ monocytes in green and Ly6C- PE^+^ monocytes in red. Green and red signals are merged in yellow. Bar graphs represent the intensity of Ly6C-PE staining (MFI) on non-clustered and clustered monocytes at 1 or 16 hrs post Ly6C-PE mAb injection across 2 replicate experiments (n=2-4 mice). *r* corresponds to the ratio of MFI between clustered and non-clustered monocytes. (B) Gating strategy to identify by flow cytometry Ly6C-PE^hi^ and Ly6C-PE^low^ monocytes, after gating on Ly6C-PerCpCy5.5^+^ monocytes following the experimental design as described above. (C) Representative dot plots and FACS histograms of cell-surface ICAM-1 and CD86 expression on Ly6C-PE^hi^ and Ly6C- PE^low^ monocytes are shown. Bar graphs pool 2 independent replicate experiments with each symbol corresponding to one mouse and indicated p-value (n=5 mice).

**Figure S7, related to Figure 7.**
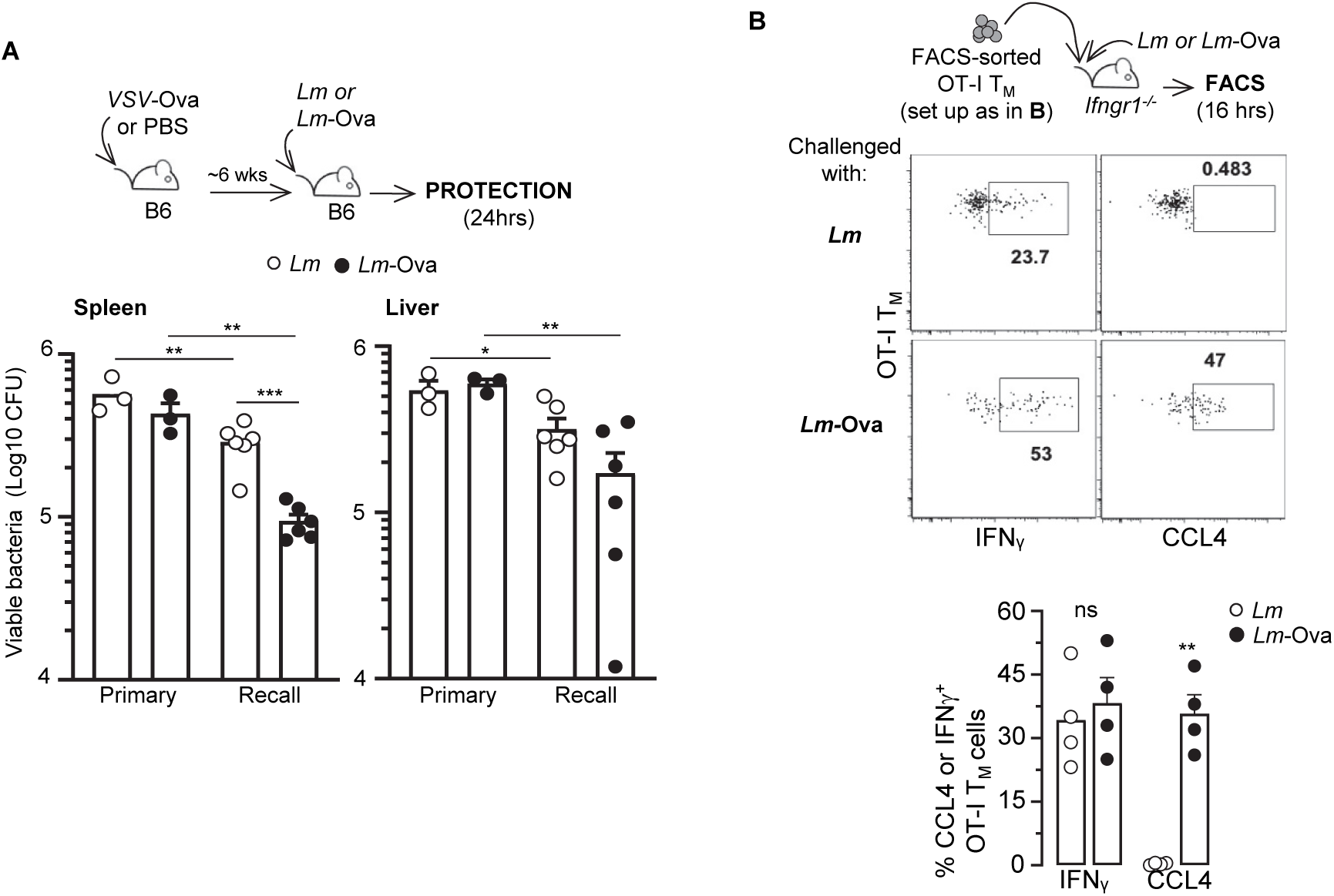
(A) Mice were immunized with 2×10^5^ PFU *VSV*-Ova or injected with PBS, and ∼6 wks later challenged with 10^6^ *Lm* or *Lm*-Ova. Spleens and livers from challenged mice were harvested 24 hrs later and *Lm* CFUs determined after plating. Bar graphs show 1 of 2 representative experiments with each symbol corresponding to 1 individual mouse. (B) 2×10^5^ OT-I T_M_ cells induced using the depicted experimental set up, were transferred in *Ifngr1^-/-^* mice, and mice were next challenged with 10^6^ *Lm* or *Lm-*Ova for 16hrs. OT-I Td^+^ T_M_ cells were stained for cell surface CD8, CD3 and intracellular CCL4 and IFNγ. Representative FACS dot plots are shown and bar graphs pool 2 independent replicate experiments (n=4 mice) with indicated p-values.

**Movie S1. Dynamic behavior of cognate antigen- *versus* inflammation-stimulated memory CD8^+^ T cells in CCR2^+^ monocyte clusters during recall infection.** Representative time-lapse movie showing cognate antigen (OT-I, red) and inflammation-stimulated gBT-I (green) CD8^+^ T_M_ cells in CCR2^+^ monocyte clusters (CCR2^CFP^, blue) at ∼16 hrs post challenge with *Lm-*Ova.

**Movie S2. Dynamic behavior of inflammation-stimulated memory CD8^+^ T cells in CCR2^+^ monocyte clusters during recall infection.** Representative time-lapse movie showing inflammation-stimulated (OT-I, red and gBT-I, green) CD8^+^ T_M_ cells in CCR2^+^ monocyte clusters (CCR2^CFP^, blue) at ∼16 hrs post challenge with *Lm*.

**Movie S3. Dynamic behavior of cognate antigen-stimulated memory CD8^+^ T cells in CCR2^+^ monocyte clusters during recall infection.** Representative time-lapse movie showing cognate antigen (OT-I, red and gBT-I, green) CD8^+^ T_M_ cells in CCR2^+^ monocyte clusters (CCR2^CFP^, blue) at ∼16 hrs post challenge with *Lm*-Ova-gB.

Table S1. List of genes up/down regulated for Ag/Infl., Infl. and Ag/Inf./Infl CD8^+^ T_M_ cells as defined in Figure 1.

Table S2. GO pathways for Fig. 1E

Table S3. Table for antibodies and other reagents

