## Supplementary figures and images for "Memory CD8^+^ T cells mediate early pathogen-specific protection through localized delivery of chemokines and IFNγ to clusters of inflammatory monocytes"

### supplemental figures

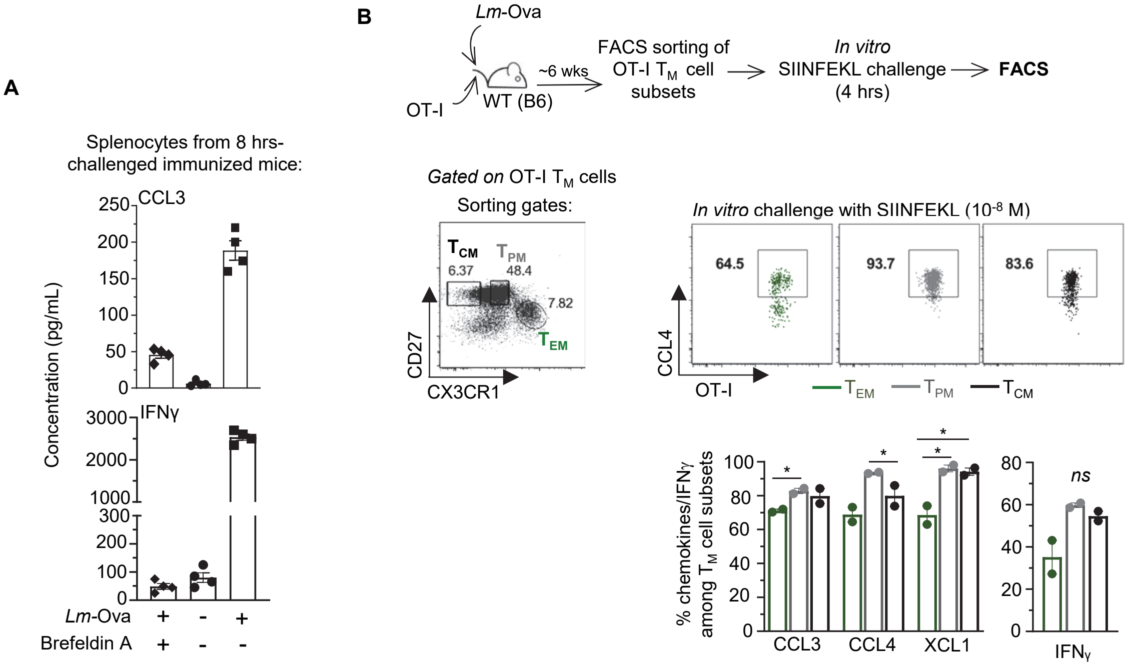

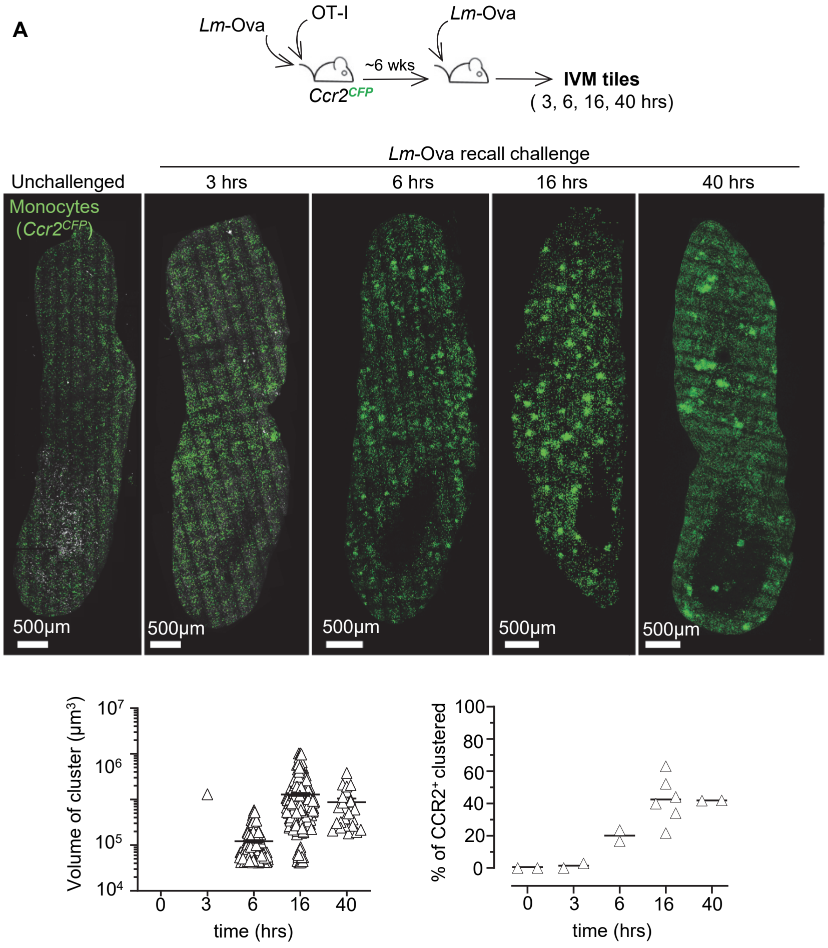

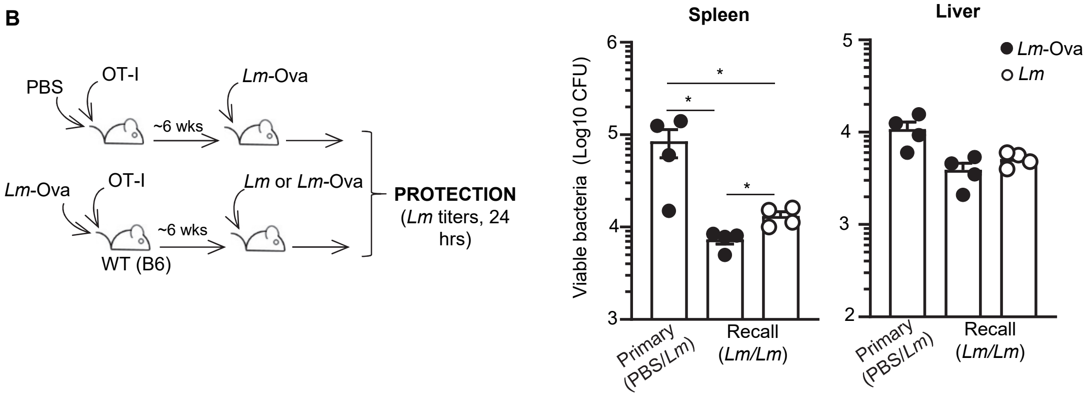

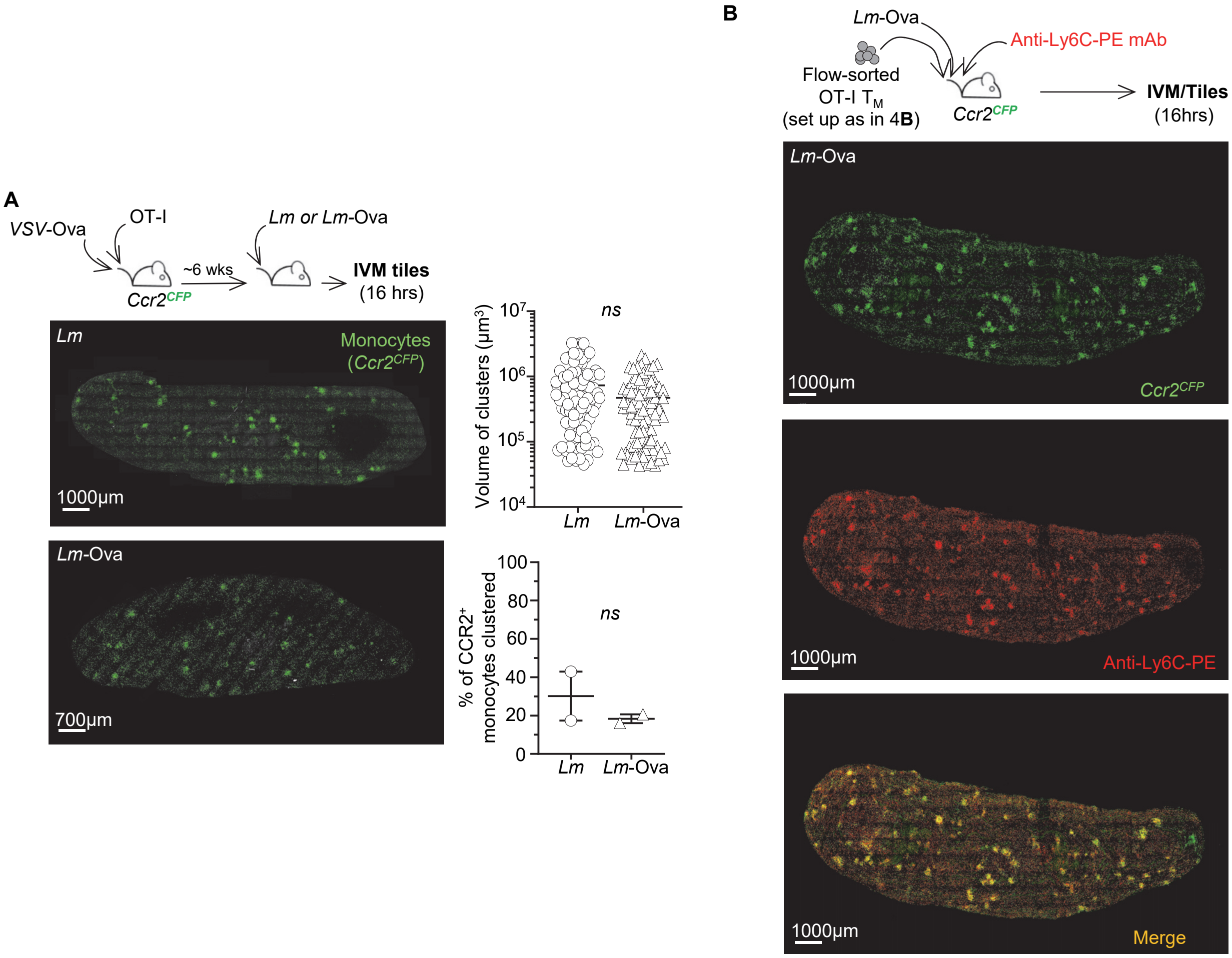

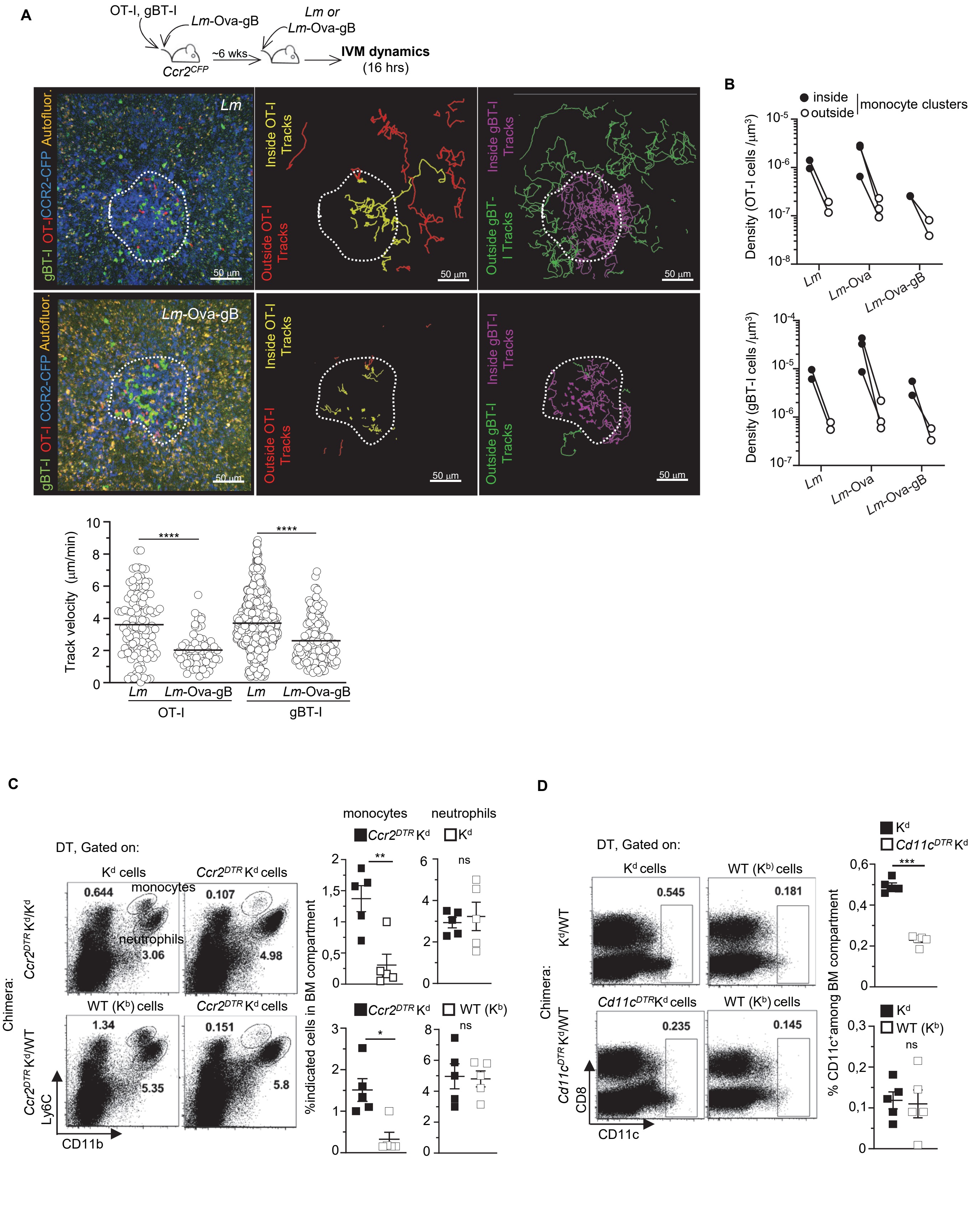

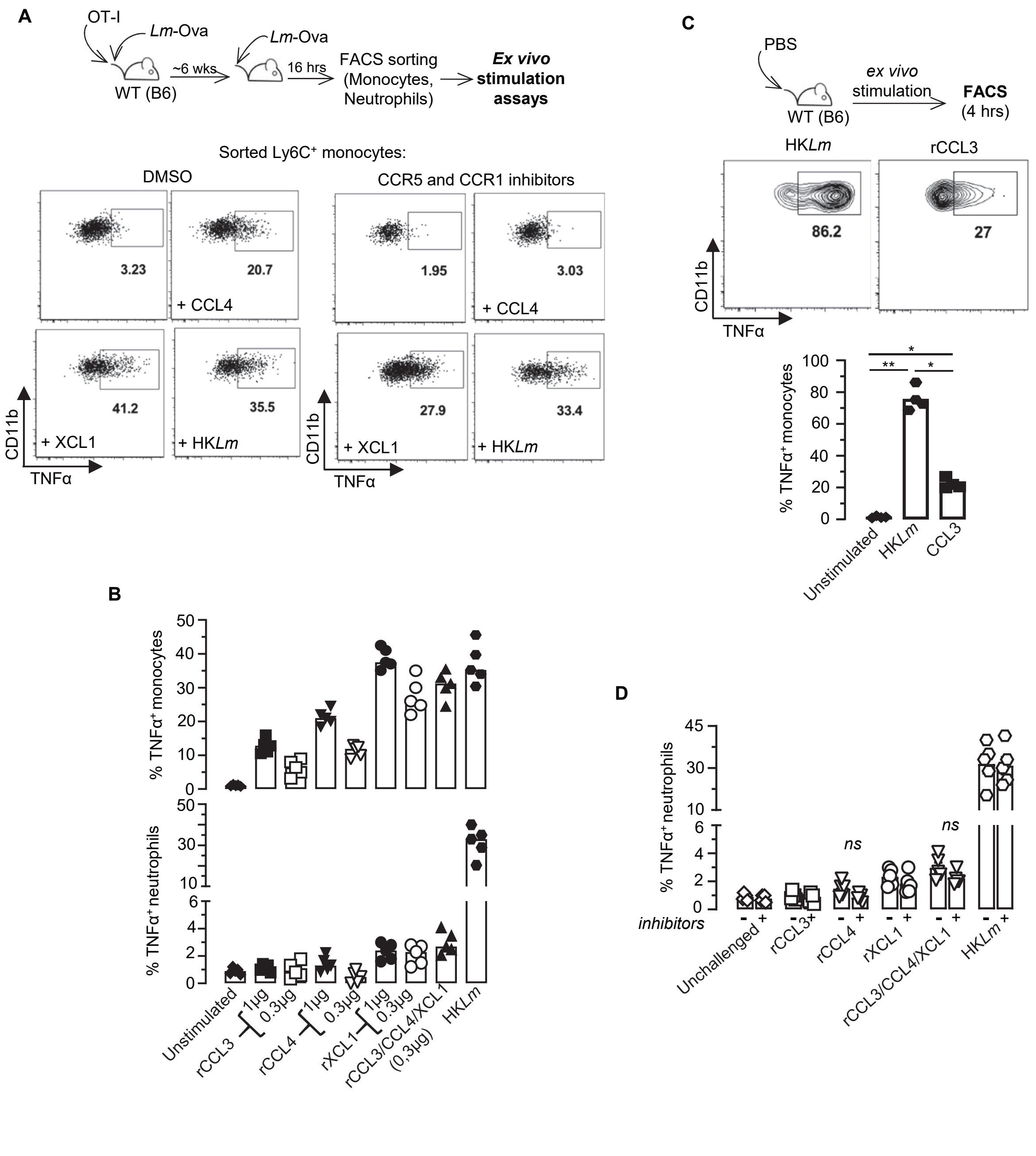

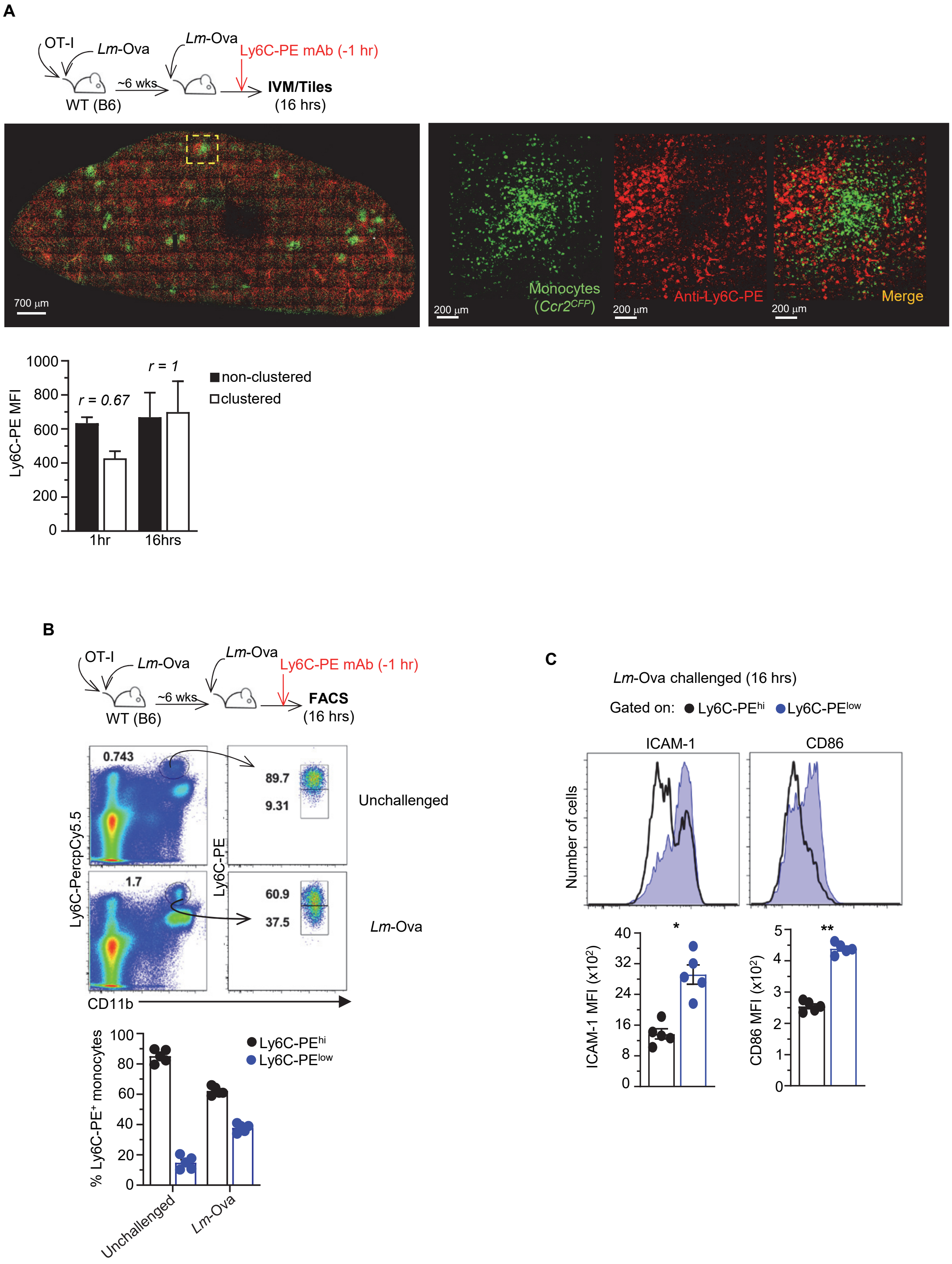

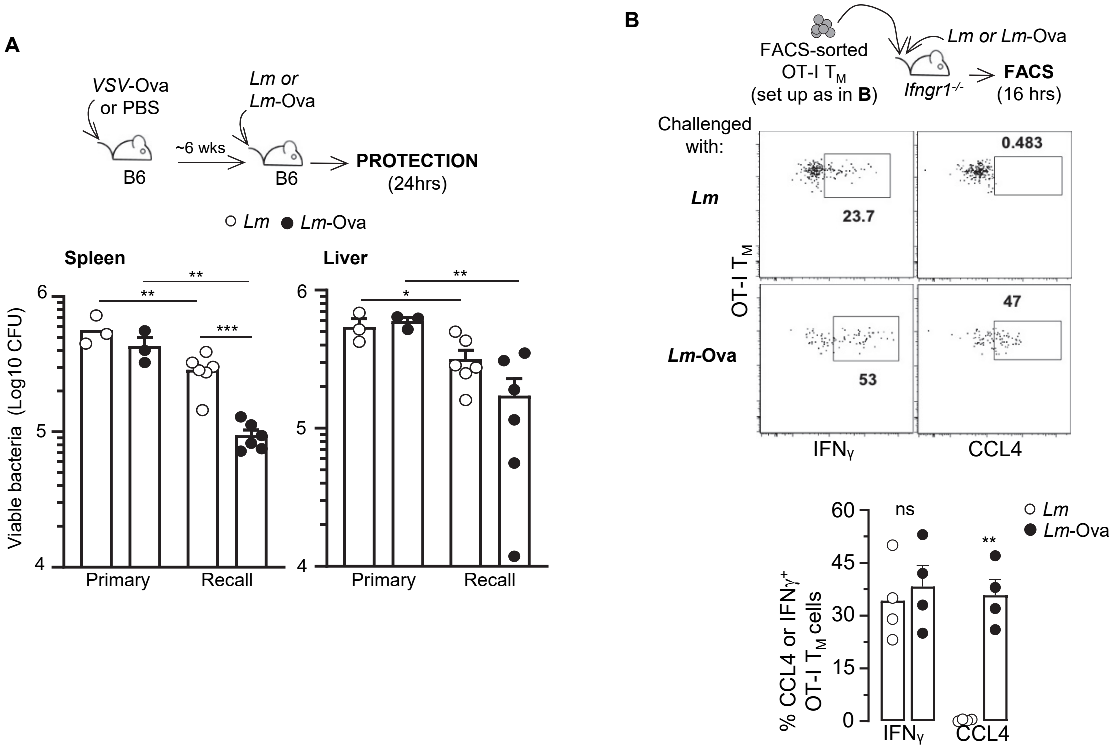
